# Under pressure: Keratin 9 regulates mechanosensitive YAP1 signaling in palmoplantar epidermis

**DOI:** 10.1101/2025.09.15.676359

**Authors:** Sarah N. Steiner, Eric Horst, Evelyn S. Chiu, Mitre Athaiya, Craig N. Johnson, Joseph Y. Shen, Michelle L. Kerns, Joseph Kirma, Rebecca L. McCarthy, Syed F.H. Shah, Ofer Sarig, Liat Samuelov, Eli Sprecher, Edel A. O’Toole, Johann Gudjonsson, Geeta Mehta, Ramiro Iglesias-Bartolome, Pierre A. Coulombe

## Abstract

The palmoplantar epidermis, in adapting to the exceptional mechanical strain it bears during locomotion, is molecularly and histologically distinct from other skin body sites. The mechanisms specifying and maintaining its unique identity remain incompletely defined. Here, we identify the type 1 keratin 9 (*KRT9*/K9), a protein uniquely expressed in the palmoplantar epidermis, as a key modulator of mechanosensitive YAP1 signaling in response to postnatal mechanical compression. K9 loss-of-function variants, as observed in human *KRT9* palmoplantar epidermal differentiation disorder (*KRT9*-pEDD), result in aberrant YAP1 subcellular partitioning and elevated levels of the stress-induced keratin 16 (*KRT16*/K16). *Krt9* null mice recapitulate these molecular phenotypes as early as postnatal day 3 (P3). We further identify dynamic, YAP1-dependent regulation of *Krt16*/K16, upstream of *Krt9*/K9 expression, during early postnatal development *in situ* and in response to mechanical compression of keratinocytes *ex vivo*, highlighting the role of mechanical stress in epidermal specification. Mechanistically, K9 interacts with the YAP1 binding protein 14-3-3σ and sequesters YAP1 in the cytoplasm, inhibiting its transcriptional activity; these functions are disrupted by *KRT9*-pEDD-causing variants in K9. Finally, we demonstrate that genetic or pharmacological inhibition of YAP1 ameliorates palmoplantar keratoderma in *Krt9* null mice. These findings reveal a role for mechanical cues in specification and maintenance of palmoplantar skin and suggest new therapeutic interventions for inherited palmoplantar epidermal differentiation disorders (pEDDs).

## INTRODUCTION

Specification of gene expression, cell fate, and tissue function is orchestrated by dynamic interplay between developmental cues and environmental factors. The epidermis offers a striking example of this complexity: as the body’s primary barrier, it mediates essential and multimodal protective functions while adapting to diverse cues and pressures ^1^. The epidermis of palmoplantar skin, located on the ventral aspect of the hands and feet, endures a substantial mechanical burden during locomotion and has unique molecular and histological adaptations ^2, 3^, among them the expression of keratins 9 (gene *KRT9*; protein K9) and 16 (*KRT16*/K16) in its differentiating layers^2^.

Keratins are an abundant and diverse family of intermediate filament (IF)-forming proteins expressed in epithelial cells across the body ^4^. The regulation of keratin protein expression is tightly choreographed depending on body site, differentiation, and context including stress ^5^. Two distinct types of keratins, I and II, interact in an obligatory fashion to yield mature intermediate filaments ^6^. Keratin 9 (*KRT9*/K9) is a type 1 keratin that is uniquely expressed in the suprabasal (differentiating) layers of human and mouse epidermis ^2, 3, 7, 8^. K9’s presumptive type II polymerization partners are keratin 1 (*KRT1*/K1) and keratin 2 (*KRT2*/K2), respectively at early and late stages of differentiation ^2, 9^. Pathogenic variants in *KRT9*/K9 cause *KRT9* palmoplantar epidermal differentiation disorder (*KRT9*-pEDD), formerly epidermolytic palmoplantar keratoderma (EPPK), an autosomal dominant disorder characterized by fragility of suprabasal, differentiating keratinocytes, acanthosis, and hyperkeratosis in palmoplantar skin ^10, 11^, suggesting a loss of mechanical integrity and defects in tissue homeostasis ^10^. *Krt9*^-/-^mice recapitulate a clear *KRT9*-pEDD-like phenotype ^12^, developing macroscopic lesions in the footpad skin by 3.5 weeks of age, tissue fragility, and hyperkeratosis. There is, as yet, no successful clinical treatment for *KRT9*-pEDD, and the molecular mechanisms underpinning the etiology of the palmoplantar lesions are not well-understood.

While *KRT9*-pEDD is, individually, a rare disease with an estimated worldwide incidence of 2.2–4.4 per 100,000 live newborns ^13^, the clinical presentation of palmoplantar keratoderma arises from pathogenic variants identified in over 50 different genes ^11^. For example, pathogenic variants in keratin 16 (*KRT16*/K16), a type I keratin associated with stress response in interfollicular epidermis ^14^, result in pachyonychia congenita (PC), a condition that entails severe, but non-epidermolytic, palmoplantar keratoderma ^15^. Incidentally, several findings have illuminated a potential link between *KRT16*/K16 and *KRT9*/K9 in palmoplantar skin; *KRT9*/K9 levels are decreased in PPK lesions of individuals with PC ^16^ and in *Krt16*^-/-^ mice ^17^, which spontaneously develop PPK-like lesions. K16, on the other hand, is strongly upregulated in *Krt9*^-/-^ mice ^12^. The dynamic interplay between these two keratins in palmoplantar skin is underappreciated and its significance unresolved.

Here, we identify a novel role of K9 in negatively modulating YAP1 signaling in human and mouse palmoplantar epidermis, suggesting a pathogenic mechanism relevant to palmoplantar keratoderma and disorders that result in it. Our findings reveal that loss of K9 function in humans through dominant-negative missense alleles disrupts YAP1 subcellular partitioning and induces upregulation of K16. *Krt9*^-/-^ mice recapitulate these molecular phenotypes, starting in early postnatal development. In development, we show that *Krt9*/K9 expression increases after a transient spike in *Krt16*/K16 in early perinatal life. This spike is YAP1-dependent, and *ex vivo* mechanical compression is sufficient to induce both *KRT9* and *KRT16* in a YAP1-dependent manner, establishing a critical link between mechanotransduction and body site-specific identity in the palmoplantar epidermis. We also show that K9 is sufficient to regulate the partitioning of YAP1 between the cytoplasm and nucleus *in vitro*, whereas *KRT9* pathogenic variants fail to do so. Crucially, we find that pharmacological or genetic inhibition of YAP1 rescues the *KRT9*-pEDD-like lesions and reduces *Krt16*/K16 expression in the *Krt9*^-/-^mouse, highlighting a prospective avenue for pharmacological treatment of *KRT9*-pEDD. By illuminating the crosstalk between mechanical cues and development, these findings provide mechanistic insight into the specification and maintenance of palmoplantar keratinocyte identity, as well as potential clinical interventions for *KRT9*-pEDD and other disease(s) manifesting with palmoplantar keratoderma.

## RESULTS

### Suprabasal nuclear YAP1 and K16 are elevated in patient KRT9-pEDD samples

Previous studies have identified mechanical fragility in the clinically-involved palmoplantar skin of *KRT9*-pEDD patients ^10^. We investigated whether *KRT9*-pEDD lesions exhibit defects in YAP1 signaling, a mechanotransducive pathway of potential relevance ^18, 19^. YAP1 has been implicated in the balance of proliferation and differentiation in keratinocytes of the interfollicular epidermis ^20, 21^. Punch biopsies obtained from lesional skin of patients clinically presenting with *KRT9*-pEDD, as well as trunk and plantar skin from healthy volunteers, were collected (see Methods) and stained via immunofluorescence (IF) for YAP1 and K9 (**Fig. 1A**). Variants in *KRT9* were confirmed (p.R163W, p.N161S, and p.M156V). Using keratin 14 (*KRT14*/K14) to mark all progenitor keratinocytes, we quantified the relative occurrence of nuclear YAP1 in basal (K14+) and suprabasal (K14-) cells (**Fig. 1B**). Consistent with previous reports in trunk epidermis ^22^ ^23^, a strong IF signal for YAP1 occurs in both the nucleus and cytoplasm in basal keratinocytes of the trunk and palmoplantar epidermis, and otherwise YAP1 occurs as a weaker and diffuse signal in the cytoplasm of suprabasal keratinocytes in healthy skin (**Fig. 1A**). A small population of K14-, nuclear YAP1+ keratinocytes occurs in the terminally differentiating layers of both neutral and plantar human epidermis (**Fig. 1A-C**). However, compared to either trunk or plantar samples from healthy volunteers, lesional skin from *KRT9*-pEDD patients exhibited strong nuclear YAP1 signal in the suprabasal epidermis (**Fig. 1A-C**), expanding the K14-, nuclear YAP1+ population by almost 10-fold. These YAP1+ nuclei occurred significantly further from the basal epidermis, as compared to either healthy trunk or plantar epidermis, correlating with the increased thickness of the K14- suprabasal layers (**Fig. 1C-D**). In keratinocytes of healthy plantar epidermis, high K9 fluorescence intensity was negatively correlated to nuclear YAP1 intensity (Pearson r = −0.2615; p=0.0015); however, in *KRT9*-pEDD palmoplantar epidermis, higher K9 expression instead positively correlated to nuclear YAP1 fluorescence intensity (Pearson r=0.2954, p=0.0016) (**Fig. 1E**). These data suggest that *KRT9*-pEDD is correlated with aberrant suprabasal nuclear YAP1.

**FIGURE 1:**
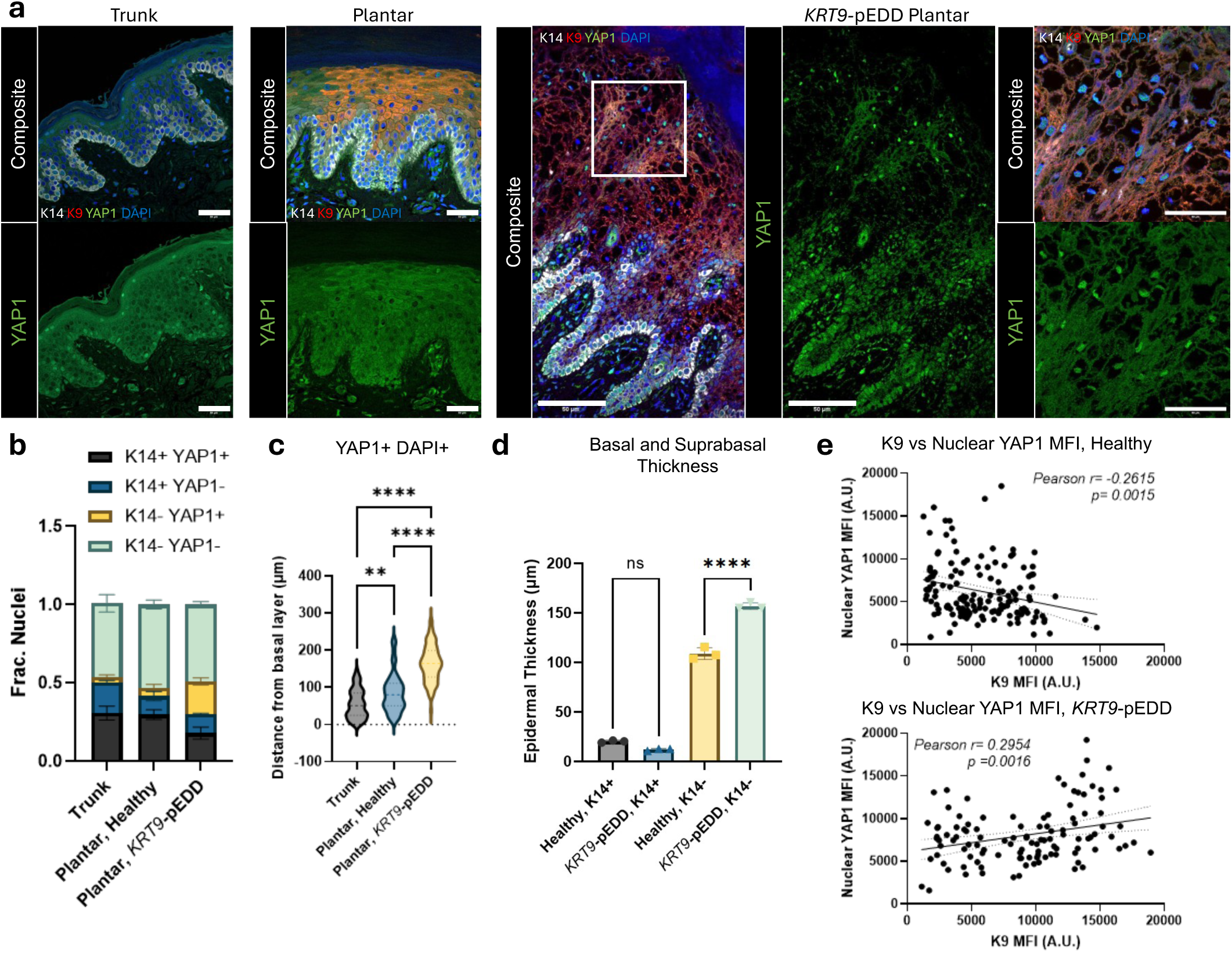
YAP1 is mislocalized in *KRT9*-pEDD patient samples. **a**) Immunofluorescence of YAP1 in human healthy trunk epidermis, healthy plantar epidermis, and the plantar epidermis of patients bearing *KRT9* variants causing *KRT9*-pEDD. *KRT9*-pEDD plantar epidermis is thicker and exhibits aberrant suprabasal nuclear YAP1. White = K14, red= K9, green =YAP1, blue=DAPI. Scale bare =50µm. **b**) Quantification of A); *KRT9*-pEDD plantar lesions exhibit a significant increase in the fraction of suprabasal K14-, nuclear YAP1+ cells. N=2 samples of trunk epidermis; N=4 healthy plantar and *KRT9*-pEDD plantar samples. 3 images per individual quantified. Bars represent mean +/- SEM. Two-way ANOVA analysis. **c**) YAP1 positive nuclei quantified as a factor of distance from basal lamina (“0”) across healthy trunk, healthy plantar skin, and *KRT9*-pEDD plantar skin. N=4 individuals/condition; one-way ANOVA analysis. **d**) Quantification of the thickness of K14+ and K14- epidermal layers in healthy and KRT9-pEDD plantar skin. N=3 individuals/condition; one-way ANOVA analysis. Dots represent individuals. Bars represent mean +- SEM. **e**) Quantification of K9 mean fluorescence intensity versus nuclear YAP1 mean fluorescence intensity in individual cells of healthy (top) and *KRT9*-pEDD (bottom) plantar epidermis. A.U.= arbitrary units. Healthy skin: Pearson r= −0.2615; p=0.0015. *KRT9*-pEDD epidermis: Pearson r= 0.2954, p=0.0016. N=4 individuals counted; approximately 120 cells total counted in each condition.

*KRT9*-pEDD patient samples also displayed an increase in K16 expression (**Suppl. Fig. 1A-B**). Unlike trunk skin, K16 is constitutively expressed in healthy plantar skin epidermis ^2, 7^. In *KRT9*-pEDD patient samples, K16 and K9 fluorescence intensity do not have any meaningful correlation, and K16 fluorescence intensity is high irrespective of the K9 level in individual cells (**Suppl. Fig. 1C**). In individual cells of the healthy palmoplantar epidermis, however, K9 and K16 staining display a significant negative correlation (**Suppl. Fig. 1C**), a finding corroborated on the transcript level through single-cell RNA-seq analysis (**Suppl. Fig 1D;** ref. ^7^). This contrasts with other differentiation-specific keratins; for instance, *KRT9* and *KRT10* exhibit a strong positive correlation (**Suppl. Fig. 1D**), as do *KRT9* and its presumptive type 2 partner *KRT1,* and *KRT16* and its type 2 partner *KRT6A* (**Suppl. Fig. 1D**). Interestingly, the top 10% of *KRT9*-expressing cells exhibit enrichment in GO terms relating to formation of the cornified envelope, programmed cell death, and keratinocyte differentiation, while the top 10% of *KRT16*-expressing cells enrich terms relating to innate immune activation and wound healing ^5^ (**Suppl. Fig. 1E**). In summary, *KRT9*-pEDD patients experience both dysregulation of YAP1 subcellular portioning and increased K16 expression. These observations suggest that K9 may negatively modulate YAP1 nuclear localization and K16 expression in healthy human plantar epidermis.

### Krt9^-/-^ mice display aberrant YAP1 localization in the suprabasal layers of the palmoplantar epidermis

To test the molecular mechanism(s) underpinning the formation of *KRT9*-pEDD keratodermas, we set out to identify potential YAP1 defects in a mouse model of *KRT9*-pEDD. *Krt9*^-/-^ mice, as related above, develop clear *KRT9*-pEDD-like manifestations by 2.5-3 weeks of age (ref. ^12^ and **Suppl. Fig 2**), and exhibit abnormally thickened palmoplantar epidermis as early as postnatal day 10 (P10) (**Suppl. Fig 2A-B**). Furthermore, we identified increased cross-sectional cellular area of keratinocytes in *Krt9*^-/-^ mice at 3.5 weeks of age (**Suppl Fig. 2C-D**). Accordingly, we focused our attention on the first 10 days of life to detect early molecular alterations. At postnatal day 0 (P0) WT and *Krt9*^-/-^ mice have similar YAP1 localization throughout the palmoplantar epidermis (**Fig. 2A, 2D**). As early as P3, however, *Krt9*^-/-^ mice exhibit aberrant nuclear-localized YAP1 in the suprabasal layers, compared to their WT littermates, as observed by immunofluorescence (**Fig. 2B, 2D**). This molecular phenotype has worsened by 3.5wks of age, when gross lesions are observable in *Krt9*^-/-^ mice (**Fig. 2C-D**). While overall levels of YAP1 are unaltered (**Fig. 2E**), we find that the ratio of phosphorylated (Ser127) YAP1 to total YAP1 is significantly attenuated in *Krt9*^-/-^ animals at P3 via Western blot (**Fig. 2E**). Phosphorylated YAP1 is the form sequestered in the cytoplasm and therefore inactive ^24^; the reduction of phospho-YAP1 in the footpad skin of *Krt9*^-/-^ mice correlates to the increase in its nuclear localization. Furthermore, the expression levels of the YAP1 binding protein 14-3-3σ are also significantly elevated in *Krt9*^-/-^ animals by P3 (**Fig. 2E**). We next examined, via bulk RNA sequencing, transcriptional patterns altered in the *Krt9*^-/-^ mice relative to their WT littermates at P0 and P3 (**Fig. 2F-G**). Notably, *Krt19*, previously identified as a YAP1 target gene ^20, 25^, was significantly upregulated in *Krt9*^-/-^ mice at both P0 and P3. In this setting, we also identified that levels of K16, which have previously been identified as elevated in the lesions of *Krt9*^-/-^ animals ^12^, are similar in WT and *Krt9*^-/-^ littermates at P0 but are elevated in *Krt9*^-/-^ animals, relative to WT, at P3 (**Suppl. Fig. 2E**). These findings mirror those in adult human *KRT9*-pEDD lesions, provide insight as to developmental onset of *KRT9*-pEDD-like phenotypes, and suggest that an increase in nuclear YAP1 may be one of the earliest molecular alterations in the etiology of *KRT9*-pEDD-like lesions.

**FIGURE 2:**
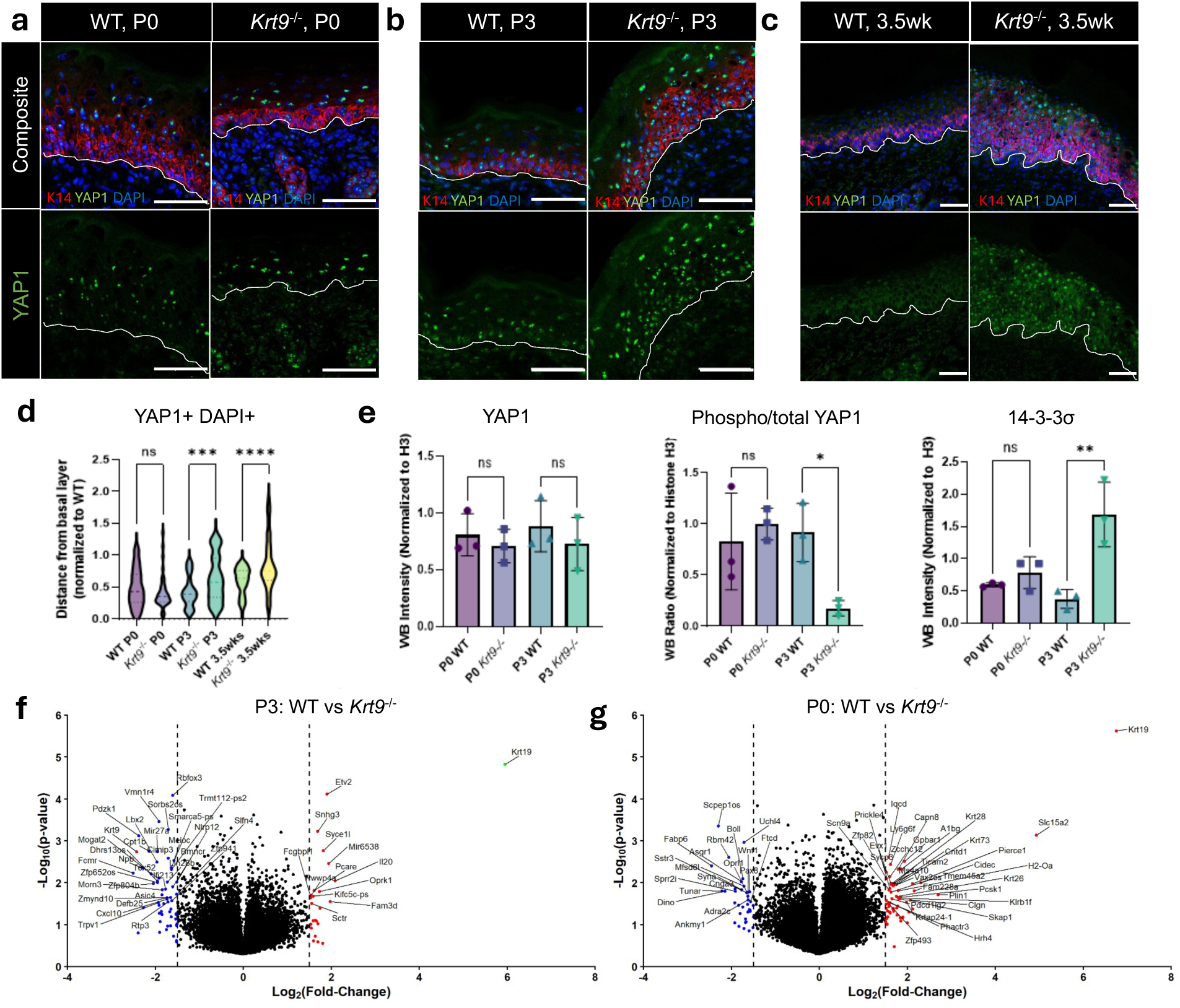
Defects in YAP1 localization in *Krt9*^-/-^ palmoplantar epidermis. a-c) Immunofluorescence of K14/YAP1/DAPI in WT/*Krt9*^-/-^ footpads. P0 (**a**), P3 (**b**), and 3.5wks (**c**) of age shown. Red=K14, green=YAP1, blue=DAPI, scale bar = 50µm. Dashed line represents border between epidermis and dermis. **d**) Quantification of YAP1 positive nuclei quantified by distance from basal layer (“0”) in (A); N=3 biological replicates per age/genotype. Normalized relative to average WT epidermal thickness at each respective age point. Data represents individual values, bars indicating mean +/- SEM. One-way ANOVA. **e)** Quantification of western blots of phospho-YAP1, total YAP1, and 14-3-3σ in WT and *Krt9*^-/-^ P0/P3 animals. Normalized to Histone H3. N=3 animals/genotype per timepoint. Data are represented as mean +/- SEM. One-way ANOVA. **f-g**) Volcano plots representing bulk RNA-seq of WT/*Krt9*^-/-^ animals at **f**) P0 and **g**) P3. N=3 per genotype. Significance cutoff at +/- 1=Log_2_(Fold Change).

### Krt9/K9 and Krt16/K16 show dynamic expression patterns in early postnatal development

Based on the early postnatal emergence of the *Krt9^-/-^*phenotype, we next examined the postnatal expression patterns of keratins in the developing palmoplantar epidermis of wildtype mice. Mice were collected at embryonic day 18.5 (E18.5), postnatal day 0 (P0), P1, P3, and P10. Whole paws were subjected to TMT-labelled quantitative mass spectrometry (TMT-MS) analysis. Upon examination of all intermediate filament species (see Methods), only five displayed dynamic regulation between E18.5 and P10: keratin 16 (*Krt16*/K16), keratin 6 isoform b (*Krt6*/K6B), keratin 17 (*Krt17*/K17), keratin 2 (*Krt2*/K2), and keratin 9 (*Krt9*/K9) (**Fig. 3A**). Notably, these dynamics include a shift from K16/K6 dominance at P0-P1 to K9/K2 dominance at P3-P10. To further examine the dynamics of these keratins during this developmental interval, whole paws were collected from WT animals at E18.5, P0, P1, P2, P3, P10, and 3.5 weeks of age and subjected to Western blot (WB). The dynamic regulation of K16 was confirmed via WB (**Suppl. Fig. 2F-G**). We further examined these dynamic patterns at the transcriptional level via bulk RNA-seq of whole paws collected from WT animals at P0 and P3. Mirroring the protein-level data, *Krt16*, *Krt6a* and *Krt6b* transcripts are enriched in P0 animals, while *Krt2* and *Krt9* transcripts are enriched in P3 samples (**Fig. 3B**).

**FIGURE 3:**
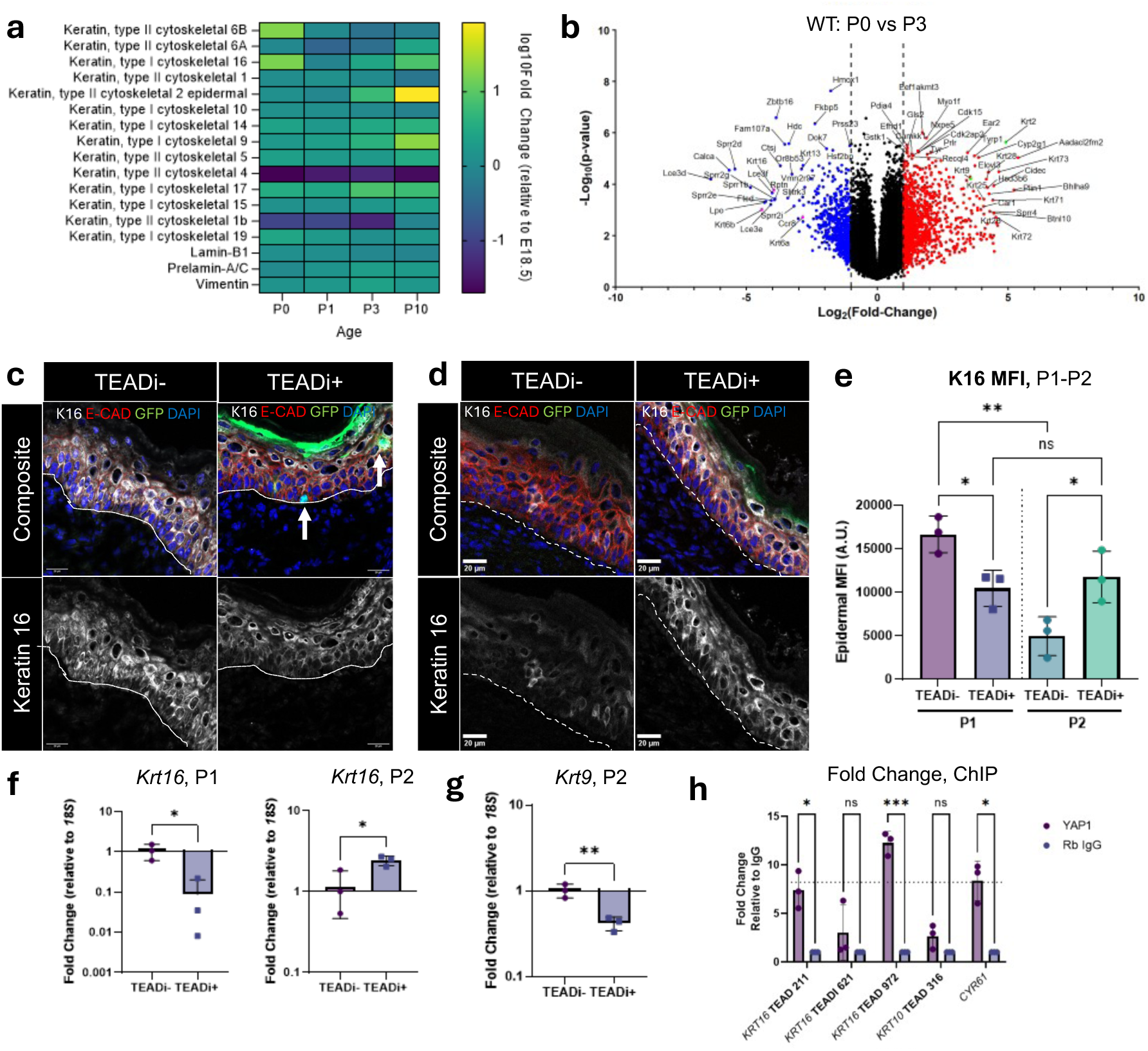
Postnatal dynamics of *Krt9*/K9 and *Krt16*/K16 are partially YAP1-dependent. **a**) Heat map of TMT 10-plex labeled quantitative MS across WT development in the paw. N=2 animals per time point. Heat map colors represent Log_10_(Fold Change) relative to E18.5; see key. **b**) Volcano plot representing bulk RNA sequencing of WT animals at P0 and P3. N=3 per timepoint/genotype. Significance cutoff at +/- 1.5=Log_2_(Fold Change) **c**) Immunofluorescence (IF) of K16 in postnatal day 1 TEADi +/- mice. White=K16, red= E-Cadherin, green= TEADi-GFP autofluorescence, representing successful recombination, blue=DAPI. Scale bar= 20µm. N=3 animals per genotype. **d**) IF of K16 in postnatal day 2 TEADi +/- mice. White=K16, red= E-Cadherin, green= TEADi-GFP autofluorescence, representing successful recombination, blue=DAPI. Scale bar= 20µm. N=3 animals per genotype. **e**) Quantification of K16 epidermal mean fluorescence intensity in **c**) and **d**); A.U.=arbitrary units. Dots represent average K16 MFI in each individual. Bars represent mean +/- SEM. **f)** *Krt16* transcript, measured via RT-qPCR, in TEADi+ mice and TEADi- littermates at P1 and P2. N=3 animals/genotype. Y-axis = Log_10_. Fold change relative to *18s*. **g**) *Krt9* transcript, measured via RT-qPCR, in TEADi+ mice and TEADi- littermates at P2. N=3 animals/genotype. Y-axis = Log_10_. Fold change relative to *18s.* **h**) ChIP-qPCR identifies two candidate YAP1 binding sites upstream of *KRT16* in hyperconfluent N/TERT2G keratinocytes. KRT16 TEAD 211, TEAD 621, and TEAD 972, as well as KRT10 TEAD 316, were examined. Results were benchmarked to signal in a known YAP1 target gene, *CYR61*, represented by the dashed line. Purple=YAP1 pulldown, blue=IgG pulldown control. Fold change relative to IgG pulldown. Each dot represents a biological replicate. Bars represent mean +/- SEM. Two-way ANOVA analysis.

To test whether the postnatal dynamics of *Krt9*/K9 and *Krt16*/K16 are YAP1-responsive in normal development, we utilized a recently described mouse line in which the expression of a synthetic TEAD inhibitor (TEADi) is placed under dual doxycycline (ROSA26 *^LSL^*^-rtTA^, ^26^) and tamoxifen (K14^CreERT^, ^27^) control in basal epidermal keratinocytes (ref. ^20^; **Suppl. Fig. 3A**). YAP1 binding to TEAD factors is essential for its transcriptional effects ^28^; hence, blocking YAP1 binding to TEAD factors should eliminate its transcriptional activity while leaving the localization, phosphorylation, and expression of endogenous YAP1 intact ^20^. TEADi+ males (K14^CreERT^ ^+/+^; ROSA26 *^LSL^*^-rtTA^ ^+/+^; *tetO*-TEADi ^+/-^) were crossed with WT C57/Bl6J females. At E17.5, pregnant females were transferred to doxycycline chow; at E18.5, IP injection of 60mg/kg tamoxifen was performed (**Suppl. Fig. 3B**). Pups were then collected at P1, when *Krt16*/K16 is transiently elevated in WT contexts, and at P2, when *Krt9* levels increase in WT palmoplantar epidermis. At P1, TEADi+ mice exhibited significantly reduced K16 staining via immunofluorescence (**Fig. 3C, 3E**), reduced K16 protein via WB (**Suppl. Fig. 3D-E**), and reduced *Krt16* transcript via RT-qPCR relative to their TEADi- littermates (**Fig. 3F**). This delay in the observed *Krt16*/K16 spike at P1 in TEADi+ perinatal mice had consequences for later *Krt9* onset; at P2, *Krt9* transcript levels were attenuated in TEADi+ mice relative to their WT littermates (**Fig. 3G**). Furthermore, at P2, *Krt16*/K16 levels were elevated in TEADi+ mice relative to their TEADi- littermates (**Fig. 3G, Suppl. Fig. 3F-G**). These data together suggest that the observed *Krt16*/K16 dynamics in early perinatal development are partially YAP1-dependent and are important for subsequent initiation of *Krt9* expression.

To examine whether YAP1-dependent regulation of *KRT16* occurs via direct binding upstream of the transcription start site (TSS), we utilized a modified N/TERT2G 3D culture protocol ^29, 30^ to generate a post-confluent, uniformly bilayered epithelium in submerged culture. Under such conditions, expression of keratinocyte differentiation markers, e.g., K10 and K16, is exclusive to the upper layer of keratinocytes ^30^. The *KRT16* locus has 3 candidate TEAD binding sites within 1kb upstream of the TSS; *KRT10* has only one (**Suppl. Fig. 3H**). ChIP-qPCR assays confirmed binding of YAP1 to two of three candidate *KRT16* TEAD sites, while it does not bind to the candidate TEAD site upstream of *KRT10* (**Fig. 3H**). Taken together, we observed that *Krt16*/K16 is transiently upregulated, in a YAP1-dependent fashion, in the postnatal palmoplantar epidermis. Delaying or reducing this YAP1-responsive increase in *Krt16*/K16 also reduces later *Krt9* onset. Based on ChIP assays, *Krt16* is a likely YAP1 target gene in the context of differentiating keratinocytes. As *Krt16*/K16 is typically considered a stress-responsive keratin ^5, 31–33^, we hypothesized that this phenomenon occurs in response to mechanical compression during the first use of the developing paw pads during nursing and early locomotion.

### Palmoplantar keratins are responsive to compressive stress in a YAP1-dependent fashion

To assess whether *KRT16* and *KRT9* are responsive to mechanical compression in keratinocyte cultures *ex vivo*, we employed a compression bioreactor described in Novak et. al. ^34^. Briefly, cells are suspended in an agarose/ collagen type 1 hydrogel and subjected to mechanical force; this 3-D hydrogel culture removes the contribution of cell-cell contacts to mechanosensing. N/TERT keratinocytes (2G subclone), selected as a substitute for human primary keratinocytes ^29, 35^, were suspended in a 1% agarose/1mg/mL collagen type I hydrogel, approximating the Young’s modulus of viable epidermis of the plantar surface ^36, 37^ and subjected to 10 kilopascals (kPa) of static compression for 24 hours. N/TERT cells were also cultured in parallel under identical conditions but without the application of compressive force (**Fig. 4A, Suppl. Fig. 4A**). To examine whether compression-induced transcriptional changes were YAP1-dependent, we further utilized the YAP1 inhibitor verteporfin (VERT). VERT has been shown to inhibit YAP1 transcriptional activity by re-localizing it to the cytoplasm, though the precise mechanism underlying this effect varies in different biological contexts ^38–40^. Inhibition efficiency was evaluated via transcription of *CYR61*, a canonical YAP1 target gene ^21, 41^. In N/TERT keratinocytes post-compression, key keratin transcripts normally expressed in the palmoplantar epidermis—*KRT9* (**Fig. 4B**), *KRT17* (**Fig. 4D**), and *KRT16* (**Fig. 4E**)—were significantly upregulated, suggesting that their expression is responsive to mechanical stress; compression-induced *KRT9*, *KRT16*, and *KRT17* were, however, absent when cells were treated with 2.5µM VERT (**Fig. 4B-E**). Similarly, *CYR61*, a well-characterized YAP1 target gene ^41^, was induced by mechanical compression and inhibited by VERT treatment (**Fig. 4C**). By contrast, *Actin*, which was upregulated in response to compression, was not significantly inhibited by VERT treatment (**Fig. 4F**), and nor was *RPS6*, a control gene that does not respond to mechanical force (**Fig. 4G**). To further test the interplay between *KRT16* and *KRT9* under mechanical compression, we employed *KRT16* knockout N/TERT keratinocytes ^42^. *KRT16* KO N/ TERT cells failed to upregulate *KRT9* while maintaining an increase in *KRT6* and *Actin* in response to compression (**Fig. 4H**).

**FIGURE 4:**
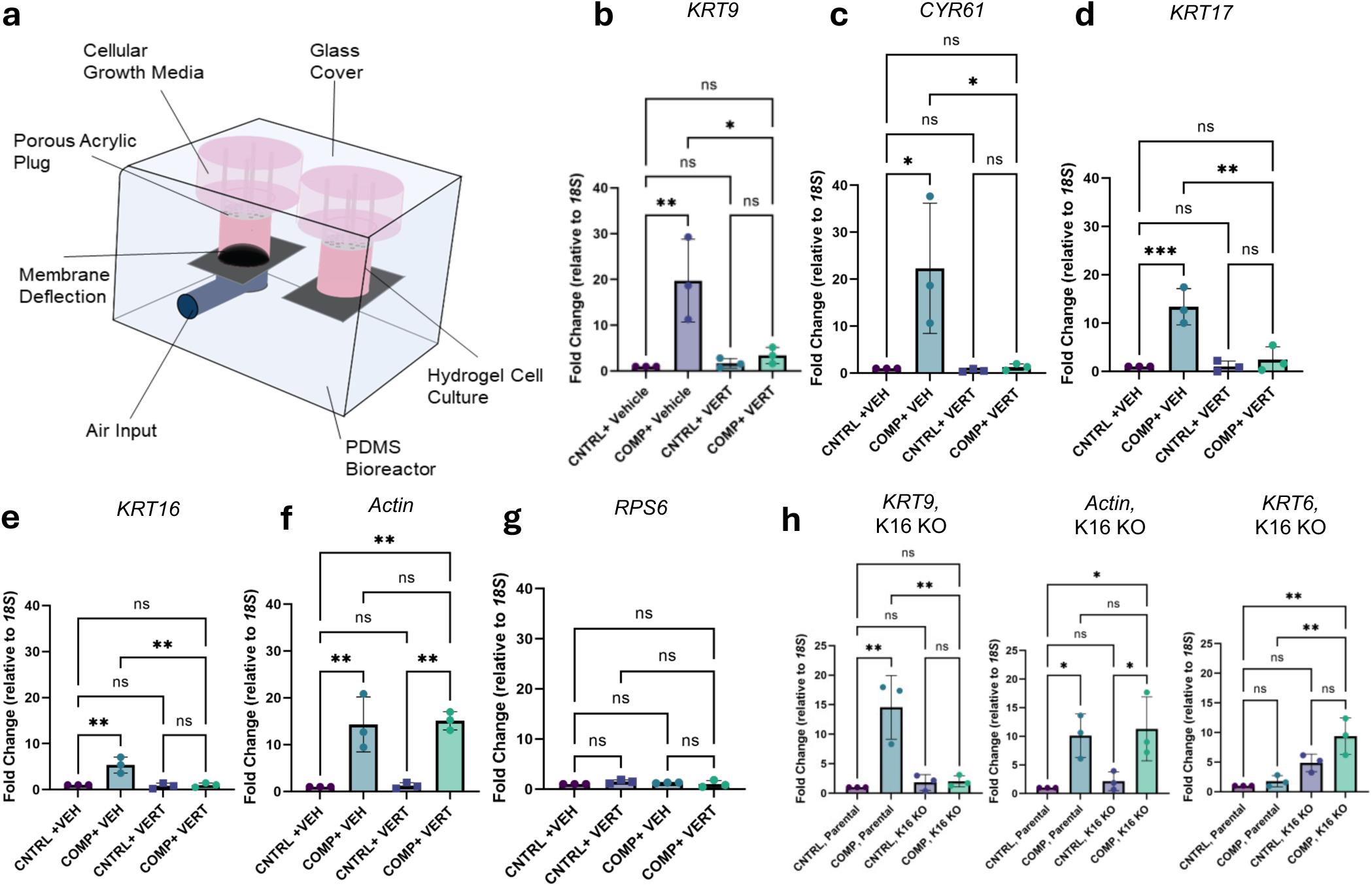
Compressive force stimulates palmoplantar keratin transcripts in a YAP1-dependent manner. **a**) Schematic of compression bioreactor, indicating components. Air is pumped into the underlying pressure chamber which deflects into the cell-laden interpenetrating hydrogel, deflecting the resistive membrane (black dome). The 3D cellular hydrogel component is held in place via a porous acrylic plug which also allows cell culture medium access from the top of this chamber. **b**) RT-qPCR of *KRT9* in vehicle and VERT-treated N/TERT2G keratinocytes subjected to compression (24hrs, 10kPa). Fold change relative to *18S*. N=3. Dark purple= Control +vehicle, light blue= compressed + vehicle, light purple= control +VERT, bright teal= compressed +VERT. **c**) RT-qPCR of *CYR61* in vehicle and VERT-treated N/TERT2G keratinocytes subjected to compression (24hrs, 10kPa). Fold change relative to *18S*. N=3. **d**) RT-qPCR of *KRT17* in vehicle and VERT-treated N/TERT2G keratinocytes subjected to compression (24hrs, 10kPa). Fold change relative to *18S*. N=3. **e**) RT-qPCR of *KRT16* in vehicle and VERT-treated N/TERT2G keratinocytes subjected to compression (24hrs, 10kPa). Fold change relative to *18S*. N=3. **f**) RT-qPCR of *Actin* in vehicle and VERT-treated N/TERT2G keratinocytes subjected to compression (24hrs, 10kPa). Fold change relative to *18S*. N=3. **g**) RT-qPCR of *RPS6* in vehicle and VERT-treated N/TERT2G keratinocytes subjected to compression (24hrs, 10kPa). Fold change relative to *18S*. N=3. **h**) RT-qPCR of *KRT9, Actin,* and KRT6 in parental and K16 KO N/TERT2G keratinocytes subjected to compression (24hrs, 10kPa). Fold change relative to *18S*. N=3. Dark purple= control parental, light purple= parental compressed, dark blue= K16 KO control, light blue= compressed K16 KO.

For additional characterization, human A431 epidermal carcinoma cells were selected for their epithelial character and the absence of both *KRT9* and *KRT16* expression at baseline. Compressed cells exhibited no change in proliferative activity as measured by Ki67 staining and otherwise remained viable in the setting of bioreactor culture (**Suppl. Fig. 4B-C**). In A431 cells, we further confirmed the YAP1-dependent response to mechanical compression in *KRT9*, *KRT16*, *KRT17*, and *CYR61* (**Suppl. Fig. 4D**), replicating our observations in N/TERT keratinocytes. We further examined whether mechanical compression impacted transcription of other genes involved in epidermal homeostasis, as well as other keratin genes (**Suppl. Fig. 4E**). No other tested keratin transcripts displayed a compression-responsive upregulation; of the other genes involved in epidermal homeostasis and differentiation, only *KLF4* and *WNT5A* displayed upregulation in post-compression A431 cells (**Suppl. Fig. 4E**). Both *WNT5A* and *KLF4* have been implicated in *KRT9* regulation *in vivo* ^43–45^. Taken together, these data suggest that palmoplantar-enriched keratins are responsive to mechanical compression in a YAP1-dependent fashion, and that *KRT9* induction under compression is dependent on *KRT16*.

### K9 regulates YAP1 subcellular partitioning and activity via 14-3-3σ interaction

We next examined whether K9 is sufficient to modulate the subcellular partitioning of YAP1 and whether *KRT9*-pEDD-causing mutations disrupt this ability. To do so, we generated two K9 missense mutant constructs: K9-GFP R163→Q (K9 R163Q), which is the most common *KRT9*-pEDD-causative pathogenic variant in humans ^10, 46^, and K9-GFP C406→A (K9 C406A), representing the de-functionalization of a key cysteine residue that, when mutated in humans, is causative for *KRT9*-pEDD ^47^ (**Suppl. Fig. 5A**). Previously, we utilized transfection-permissive HeLa cells to show that K14 mediates the cytoplasmic retention of YAP1 ^48^. HeLa cells are particularly well-suited for these studies due to the strong nuclear YAP1 signal they exhibit when cultured on glass (**Fig. 5A**) and their robust expression of *CYR61*, allowing for the measurement of YAP1 subcellular localization and transcriptional activity. We transfected the mutant K9 constructs, WT K9-GFP, WT EGFP-K14 (as a positive control), and GFP alone (negative control) into HeLa cells and assessed the subcellular localization of YAP1 and the levels of *CYR61* in transfected cultures. In cells transfected with either WT K9-GFP or WT EGFP-K14, we observed a significant reduction of nuclear-to-cytoplasmic YAP1 fluorescence signal intensity ^49^, along with a concurrent reduction of YAP1 transcriptional activity (**Fig. 5A-C**). By contrast, cells transfected with either K9 R163Q-GFP or K9 C406A-GFP did not deplete YAP1 from the nucleus and did not exhibit reduced levels of *CYR61* relative to baseline (**Fig. 5A-C**). Overexpressed WT EGFP-K16 also did not result in YAP1 enrichment in the cytoplasm, despite sharing the stutter cysteine residue with both K14 and K9 (**Fig. 5, Suppl. Fig. 5A**; see Discussion).

**FIGURE 5:**
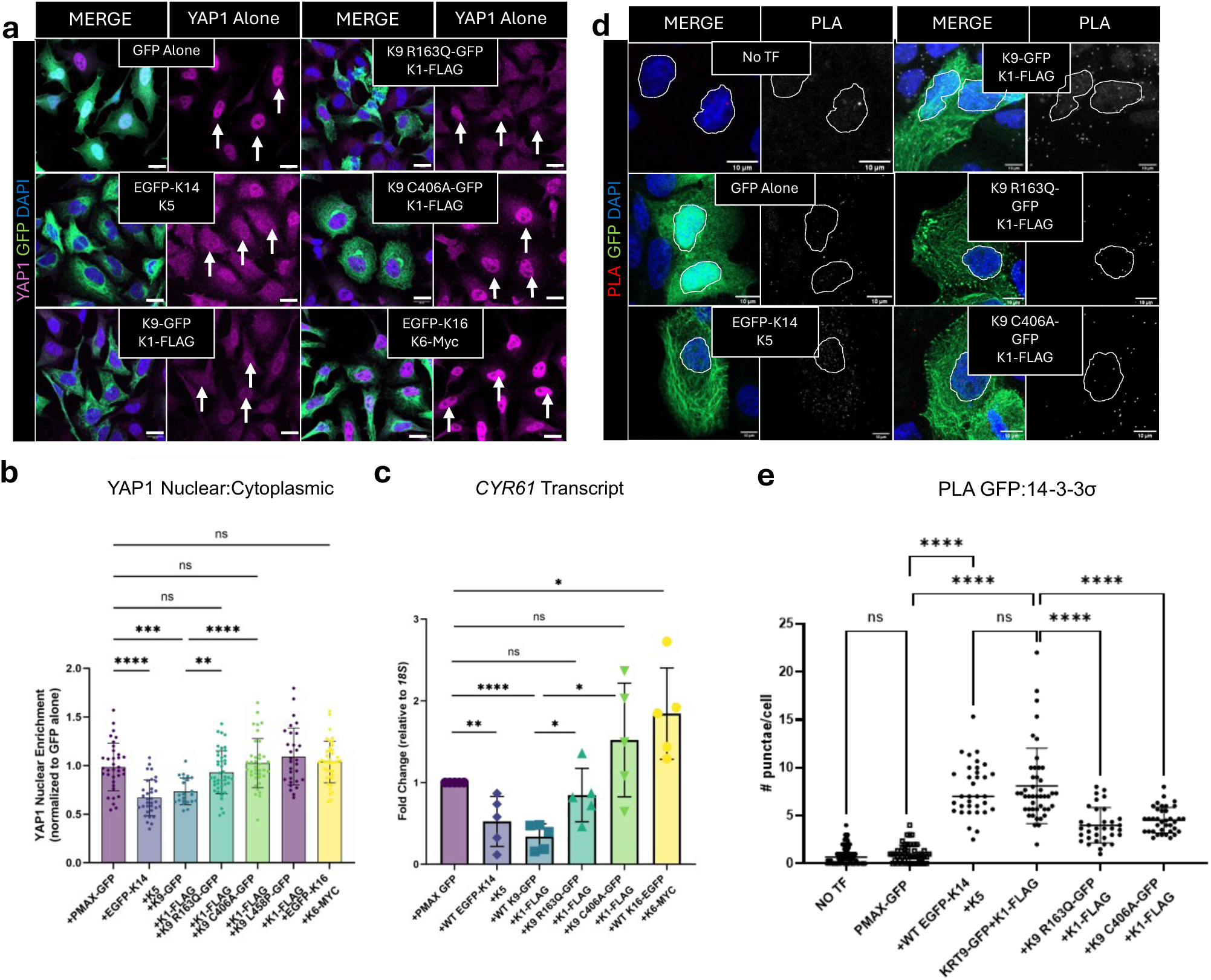
WT K9 regulates the YAP/14-3-3σ complex, while pathogenic *KRT9* variants disrupt this role. **a**) Composite IF images of YAP1 in HeLa cells transfected with WT and pathogenic K9-GFP constructs. Blue= DAPI, purple =YAP1, green = GFP autofluorescence. Scale bar =20µm. **b**) Quantification of nuclear to cytoplasmic ratio of YAP1 in (**a**). DAPI was used to define nuclear boundaries; GFP autofluorescence of transfected constructs defined cytoplasmic boundaries. N=3 biological replicates, minimum 50 cells/condition/replicate. Data represents individual cell values, bars indicating mean +/- SEM. One-way ANOVA. **c**) *CYR61* transcription in transfected HeLa cells as measured by RT-qPCR. N=5 biological replicates. Student t-test. **d**) Confocal images of proximity ligation assay (PLA) between GFP-tagged WT and *KRT9*-pEDD mimic constructs and endogenous 14-3-3σ in A431 cells. Blue=DAPI, green=GFP-tagged construct autofluorescence, red=PLA. Scale bar =10um. **e**) Quantification of PLA punctae/cell in (**d**). N=3 biological replicates; minimum 50 cells per condition per replicate. Data represents individual values, bars indicating mean +/- SEM. One-way ANOVA.

Like most cytoplasmic partners that anchor YAP1 in the cytoplasm (e.g., catenins ^22, 50^), keratin proteins are unlikely to regulate YAP1 via direct binding. Instead, cytoplasmic anchors such as catenins or K14 rely on 14-3-3 family isoforms, which bind both phosphorylated YAP1 and a phosphorylated residue on the cytoplasmic partner to form a ternary complex ^22, 48^. Phosphorylation of keratins is well-known to modulate their organization, solubility, and interactions with other signaling effectors ^51–54^. One potential interactor with phosphorylated keratins is 14-3-3σ, a 14-3-3 family protein enriched in stratified epithelia ^23^ and previously shown to promote keratinocyte differentiation in epidermis ^23^. To test whether K9 interacts with 14-3-3σ, we performed co-immunoprecipitation (co-IP) in HeLa cells co-transfected with HA-tagged 14-3-3σ and WT K9-GFP (**Suppl. Fig. 5B**). Upon HA pulldown, we observed co-IP of both WT K9-GFP and of EGFP-K14, a known 14-3-3σ interactor ^48^, indicating that these proteins indeed physically interact. We utilized computational methods predicting the likelihood of 14-3-3 binding, incorporating documented phosphorylation sites ^55^, to examine where 14-3-3 proteins are likely to bind on K9 (**Suppl. Fig. 5C**). Unlike other keratins, where the most likely 14-3-3 binding sites are phosphoserines in the head domain ^53, 56, 57^, the most highly scored residues on human and mouse K9 occur at phosphoserines in coil 1 (**Suppl. Fig. 5C**). Importantly, co-transfection of WT, K9 R163Q, or K9 C406A and its type 2 partner K1 in HeLa cells did not change overall protein levels of either YAP1 or 14-3-3σ (**Suppl. Fig. 5D-E**), suggesting that the observed changes in YAP1 transcriptional activity and nuclear-to-cytoplasmic ratios are in fact due to alterations in subcellular partitioning. Transfection with tagged keratin constructs also did not alter the nuclear or cellular area of HeLa cells (**Suppl. Fig. 5F**).

To assess whether the interaction between K9 and 14-3-3σ holds in other cell culture systems and whether it is impacted by *KRT9*-pEDD-causing pathogenic variants, we further performed proximity ligation assay (PLA) between transfected keratin constructs and endogenous 14-3-3σ in A431 cells (**Fig. 5D-E**). A431 cells expressing WT K9-GFP showed an average of 7.7 PLA+/-2.1 puncta per cell, a level comparable with cells expressing known 14-3-3σ interactor WT EGFP-K14 (**Fig. 5D-E**). By comparison, cells expressing either the K9 R163Q (3.9+/-1.8 PLA puncta/cell) or K9 C406A (4.5+/-1.2 PLA puncta/cell) variants showed significantly reduced instances of physical proximity with 14-3-3σ (**Fig. 5D-E**). Transfection of WT or mutant K9 did not affect total 14-3-3σ protein levels, observed via WB (**Suppl. Fig. 5D**), suggesting that these data result not from reduced 14-3-3σ levels but from reduced physical proximity. These data show that the property of cytoplasmic sequestration of YAP1 is conserved in K14 and K9 but not in K16, and that the introduction of pathogenic variants in K9 disrupts its ability to form a ternary complex with 14-3-3σ/YAP1 and thereby negatively regulate the transcriptional activity of YAP1.

### Pharmacological inhibition of YAP1 rescues K9-associated pathogenic phenotypes

We next hypothesized that inhibition of YAP1 may normalize the *KRT9*-pEDD-like lesions in the footpad skin epidermis of *Krt9*^-/-^ mice. To that end, we first employed a pharmacological approach using verteporfin (VERT). We tested a range of concentrations in HeLa cells in culture and found that treatment 2.5µM VERT effectively and significantly reduces the nuclear-to-cytoplasmic ratio of YAP1 (**Suppl. Fig. 6A-B**). VERT treatment restored the cytoplasmic localization of YAP1 in cells transfected with K9 R163Q and K9 C406A (**Suppl. Fig. 6C-E**). By contrast, no enhancement of YAP1 cytoplasmic sequestration was observed in cells transfected with WT EGFP-K14 or WT K9-GFP after VERT treatment (**Suppl. Fig. 6C-E**). VERT treatment of HeLa cells concurrently increases the phosphorylated-to-total YAP1 ratio (**Suppl. Fig. 6F-G**), while the steady-state levels of total YAP1 or 14-3-3σ proteins remained normal under these treatment conditions (**Suppl. Fig. 6F-G**). In this *in vitro* setting, these data suggest that VERT inhibits YAP1 primarily via impacting its phosphorylation state and therefore its subcellular partitioning.

We next tested whether VERT treatment ameliorates the *KRT9*-pEDD-like skin lesions characteristic of *Krt9*^-/-^mice. WT and *Krt9*^-/-^ mice at 3.5 weeks of age—the age of first presentation of macroscopic lesions in the front paws in *Krt9*^-/-^ mice—were treated for 7 days with vehicle on the left paw and 3mg/mL verteporfin (ref. ^58, 59^) on the right paw (**Fig. 6A**). Compared to vehicle treatment, VERT application resulted in a striking reduction (40 +/-18.9%, P<0.001) of the thickness of the living layers of the epidermis in *Krt9*^-/-^ footpad skin (**Fig. 6B-C**). To examine the mechanisms by which VERT may be normalizing the thickness of the *Krt9*^-/-^ palmoplantar epidermis, we performed RT-qPCR comparing WT and *Krt9*^-/-^ paws treated with vehicle control or VERT for 7 days and found that VERT-treated *Krt9*^-/-^ paws show a specific and significant reduction in *Krt16* (**Fig. 6D**). This pattern was corroborated on the protein level via WB and IF (**Suppl. Fig. 7A-C**). To examine whether VERT treatment impacted YAP1 localization, we performed immunofluorescence for YAP1 in WT and *Krt9*^-/-^ paws treated with vehicle or VERT (**Fig. 6E**). VERT treatment reduced the fraction of YAP1+ nuclei as a proportion of total epidermal nuclei in the treated *Krt9*^-/-^ palmoplantar epidermis to baseline levels observed in WT mice (**Fig. 6F**). Furthermore, YAP1+ nuclei occurred closer to the basement membrane, partially normalizing their distribution relative to WT (**Fig. 6F**). We further examined the levels of phospho-YAP1 in VERT-treated WT and *Krt9*^-/-^ paws via WB and found that treatment with VERT increased the ratio of phospho-to-total YAP1 in the *Krt9*^-/-^ animals, specifically (**Fig. 6G**).

**FIGURE 6:**
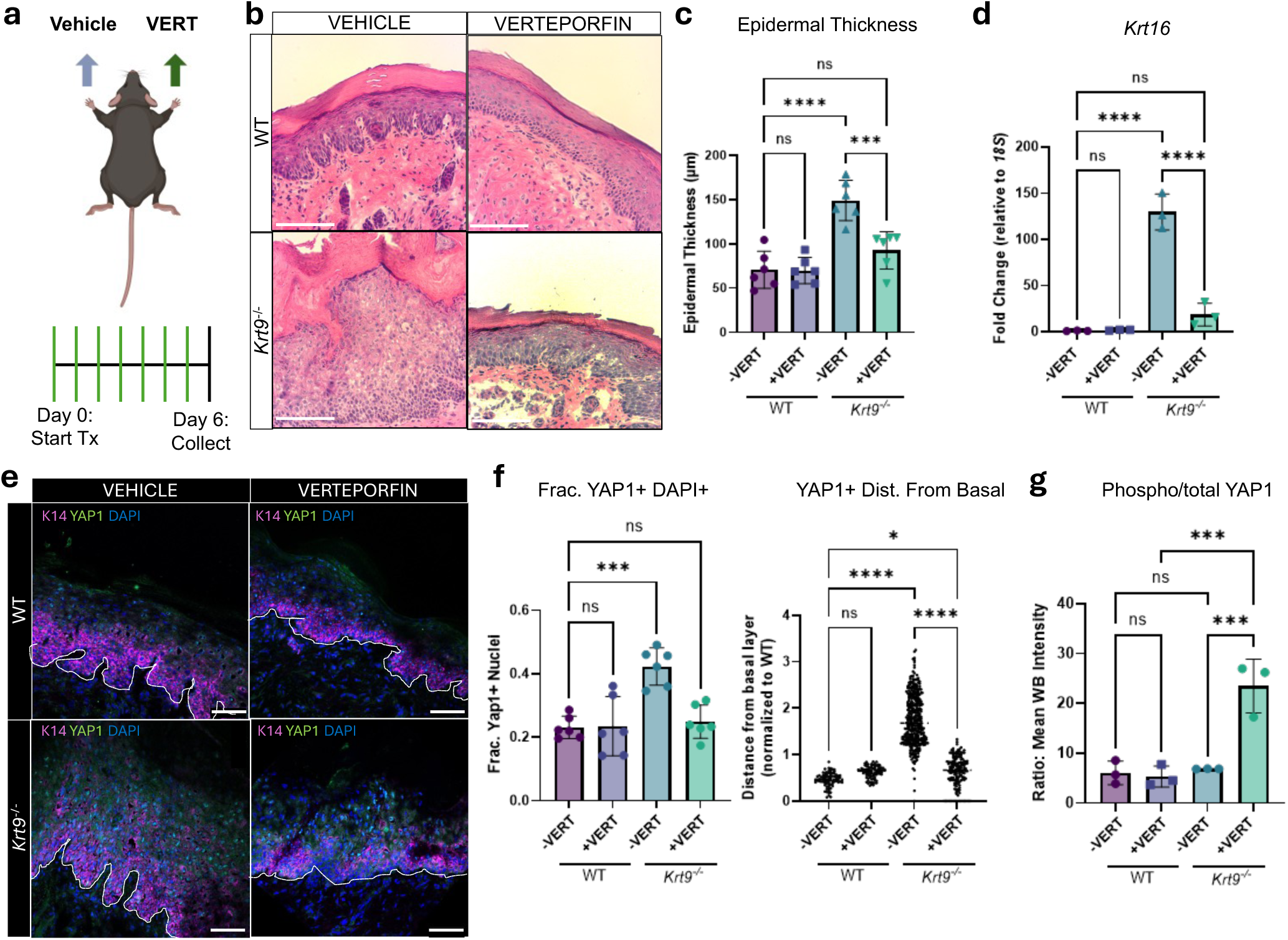
*In vivo* treatment with verteporfin rescues YAP1 localization and normalizes epidermal thickness. **a**) Schematic of verteporfin (VERT) topical treatment. 3mg/mL VERT was applied for 7 days to the right paw; vehicle (VEH) was applied to the left. **b**) H&E of vehicle and verteporfin-treated footpads of WT and *Krt9*^-/-^littermates. N= 6 animals per genotype (3 male, 3 female). Scale=100µm. **c**) Quantification of the thickness of the living layers of the epidermis in vehicle/verteporfin-treated palmoplantar epidermis. N=6 animals per genotype. Data represented as mean +/- SEM. One-way ANOVA. **d**) *Krt16* transcript, measured via RT-qPCR. Fold change relative to *18S.* N=3 animals/genotype. Data represented as mean +/- SEM. One-way ANOVA. **e**) YAP1 staining, visualized via IF, in vehicle and verteporfin-treated WT and *Krt9*^-/-^ paws. Dashed line represents boundary between epidermis and dermis. Purple=K14, green=YAP1, blue=DAPI. Scale bar=50µm. N=6 animals/genotype. **f**) Quantification of YAP1+ nuclei as a fraction of total epidermal nuclei in (**e**), Each dot represents the average fraction of YAP1+ nuclei in one mouse. N=6, minimum of 3 images per mouse. Quantification of YAP1+ nuclei with respect to their distance from the basal layer (“0”) in (**e**). Data represents individual cells, bars indicating mean +/- SEM. One-way ANOVA. **g**) Quantification of WB of phospho-to-total YAP1 ratio in WT and *Krt9*^-/-^ mice treated with vehicle or VERT. N=3 per genotype. WB normalized relative to Histone H3. Data represented as mean +/- SEM. One-way ANOVA.

YAP1 and the pro-differentiation transcription factor KLF4 ^60, 61^ have been reported to interact and oppose one another in epidermal differentiation ^20^. Interestingly, WT animals treated with VERT experience an increase in *Krt9* transcription (**Suppl. Fig. 7D**), suggesting a negative relationship between YAP1 activity and *Krt9*/K9 expression. Given previous reports that *Krt9* is a KLF4 target gene ^43^, we further examined KLF4 localization in WT and *Krt9*^-/-^ mice treated with VERT. The lesional palmoplantar epidermis of vehicle-treated *Krt9*^-/-^ mice exhibited a significantly increased fraction of KLF4+ nuclei, relative to their WT littermates, while *Krt9*^-/-^ animals treated with VERT experienced a normalization of the fraction of KLF4+ nuclei (**Suppl. Fig. 7E-F**). In the setting of WT and *Krt9*^-/-^ mice treated with VERT, these differences are likely not due to YAP1’s direct effect on the proliferative capacity of epidermal keratinocytes, as determined via Ki67 staining; aberrant nuclear YAP1 occurs primarily in the suprabasal layers, while Ki67 staining remains confined to the basal layers (**Suppl. Fig. 7G-I**). These data suggest that dysregulation of YAP1 subcellular partitioning in the *Krt9*^-/-^ mice is causative for formation of *KRT9*-pEDD-like lesions, and that pharmacological inhibition of YAP1 is a promising avenue to resolve these palmoplantar keratodermas.

### Genetic inhibition of YAP1 attenuates palmoplantar lesions in Krt9^-/-^ mice

To determine whether the beneficial impact of VERT topical treatment on the footpad skin lesions of *Krt9*^-/-^ mice reflects a specific decrease in YAP1 transcriptional activity, we crossed *Krt9*^-/-^ mice with the K14^CreERT^; ROSA26 *^LSL^*^-rtTA^; *tetO*-TEADi (TEADi+) mice described above (**Fig. 3D**, **Suppl. Fig. 3A**). The resulting *Krt9*^-/-^TEADi+ mice, along with key control mice (henceforth *Krt9*^+/+^ TEADi+ and TEADi-, *Krt9*^-/-^ TEADi-) were treated with a combination of doxycycline chow and topically applied tamoxifen (TAM) or vehicle control (Veh) to the front paws (**Suppl. Fig. 3B**). 3 days post-topical application of TAM, robust recombination was noted in animals carrying the TEADi cassette, as quantified via epidermal GFP fluorescence signal (**Suppl. Fig. 3C**); greater GFP epidermal intensity was noted in the *Krt9*^-/-^ TEADi+ mice, relative to *Krt9*^+/+^ TEADi littermates, which may be due to the thicker epidermis and longer transit time(s) required for differentiating cells to traverse it. The thickness of the living layers of palmoplantar lesions in *Krt9*^-/-^ TEADi+ animals treated with doxycycline and tamoxifen were restored to the thickness of *Krt9*^+/+^ TEADi+ and TEADi- littermates (**Fig. 7A-B**). *Krt9*^-/-^ TEADi+ mice also exhibited significantly reduced staining for K16 (**Fig. 7C-D**), corroborating the results from VERT-treated animals (**Fig. 6**). These data implicate *Krt16* as a transcriptional target of YAP1 in suprabasal keratinocytes and suggest that inhibition of YAP1 transcriptional activity is sufficient to ameliorate *Krt9*^-/-^ lesions.

**FIGURE 7:**
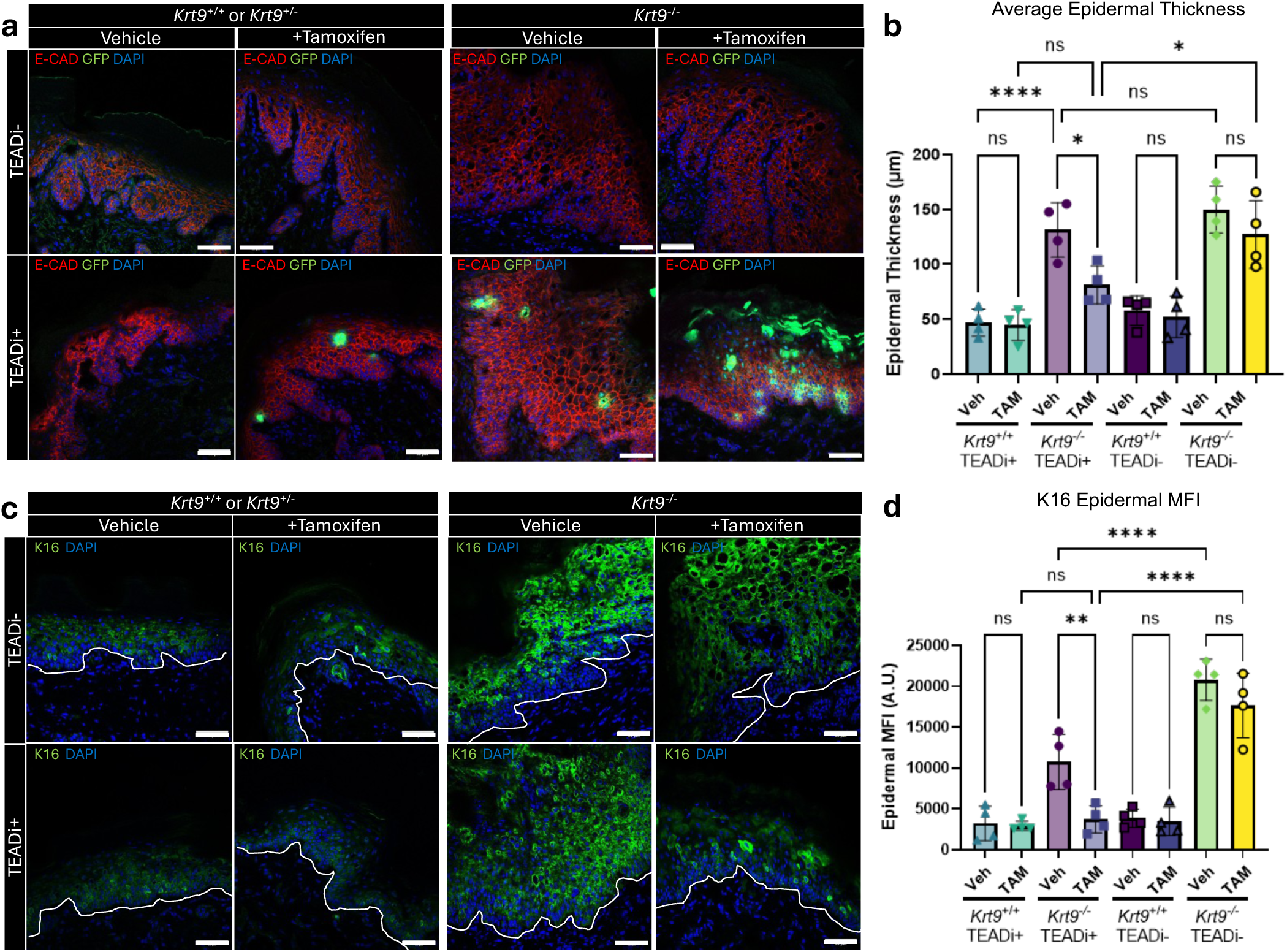
Genetic inhibition of YAP1 ameliorates *Krt9*^-/-^ lesional presentation.*\*. **a**) Composite IF images of tissue sections from *Krt9*^-/-^ TEADi+ mice and littermate controls. All mice were treated with doxycycline chow. Vehicle= left foot topically applied with vehicle, +Tamoxifen indicates right foot topically applied with tamoxifen. Red= E-cadherin, green=GFP, blue= DAPI. Scale bar=50µm. **b**) Quantification of living epidermal thickness in **a**). Minimum n=40 measurements/animal, N=4 animals/genotype/condition. Each dot represents the average in an individual mouse. Data represented as mean +/- SEM. One-way ANOVA analysis. **c**) IF of K16 (green) and DAPI (blue) in *Krt9*^-/-^ TEADi+ animals and littermate controls. Scale bar =50µm. **d**) Quantification of K16 epidermal mean fluorescence intensity (MFI) in B). N=4 animals/genotype/condition. Each dot represents the average in an individual mouse. Data represented as mean +/- SEM. One-way ANOVA analysis.

## DISCUSSION

In this study, we uncover a novel signaling function for K9 in safeguarding homeostasis during intense mechanical stress in the palmoplantar epidermis by driving the cytoplasmic sequestration of YAP1, an important mechanotransducive transcriptional effector. We illuminate the YAP1-dependent postnatal regulation of *Krt9*/K9 and *Krt16*/K16 in this specialized skin site, dynamically responding to mechanical compression. Mechanistically, we find that K9 forms a complex with 14-3-3σ, an established YAP1 binding protein, to sequester and transcriptionally inactivate YAP1. Our findings establish a causative link between loss of proper YAP1 regulation and hyperkeratosis of the palmoplantar skin, suggesting a therapeutic target for *KRT9*-pEDD and other disorders of palmoplantar epidermal differentiation.

While *KRT9*-pEDD is an individually rare condition, the clinical presentation of palmoplantar keratoderma collectively is not. Indeed, variants in more than 50 genes result in disorders of epidermal differentiation and homeostasis in palmoplantar epidermis ^11^. Accordingly, a more robust understanding of how homeostasis is established—and disrupted— in this specialized epidermal site stands to provide insight with potential clinical relevance for a wide variety of disorders. Our findings reveal a novel aspect of palmoplantar specialization: the role of mechanical stress over developmental time. *Krt9*/K9’s temporal dynamics in the developing palmoplantar epidermis distinguish it from related members of the keratin family. We find that, unlike *Krt1* ^62^ (K9’s presumptive assembly partner) or *Krt10*, *Krt9*/K9 expression does not become robust until a few days postnatally, correlating with dramatic changes to mechanical demands placed on palmoplantar skin. The differentiation-, stress-, and developmental-dependent co-regulation of type I and type II keratin *genes* as pairs (e.g., *Krt1/Krt10*) is a prominent feature in most other contexts in the epidermis ^5, 63^, marking *Krt9*/K9 as a uniquely dynamically expressed keratin. Keratin proteins, including K9, are widely appreciated for their contribution to mechanical resilience ^12, 64, 65^, and are also known to impact nuclear mechanosensing via interaction with, and force transmission through, plectins ^66^. The living layers of the palmoplantar epidermis possess remarkably different mechanical properties relative to neutral epidermis ^36^. While our results suggest that *Krt9*/K9 expression may be a response to mechanical stress, whether and how the K9 protein conveys increased mechanical resilience to keratinocytes, beyond what K10, for instance, is capable of, remains to be explored.

Previous studies on the development and specification of palmoplantar skin have identified various factors involved in *KRT9*/K9 regulation, particularly secreted factors modulating palmoplantar identity in the epidermis and underlying mesenchyme. *Ex vivo* coculture of non-palmoplantar keratinocytes with palmoplantar fibroblasts highlighted the ability of mesenchymal cues to induce *KRT9* expression ^8, 67, 68^. Fibroblasts in the distal limb exhibit a unique HOX code that is instrumental in enabling the secretion of WNT5A ^45^, which is sufficient to induce *KRT9* expression in non-palmoplantar human keratinocytes ^8, 69^. Furthermore, keratinocyte-intrinsic factors play a key part in modulating palmoplantar epidermal differentiation and expression of *KRT9*/K9. For example, variants in *WNT10A* result in loss of *KRT9*/K9 expression and the manifestation of PPK, caused in part by hypoactivation of the transcription factor KLF4 in the differentiating layers of the palmoplantar epidermis ^43^. Such factors most likely mediate the competency of the palmoplantar epidermis to express *Krt9*/K9 in response to mechanical force over developmental time. The situation differs in interfollicular epidermis, where these mesenchymal cues are absent and mechanical compression is unlikely to initiate *Krt9*/K9 expression.

In the postnatal period, we identify an important contributor to robust *Krt9*/K9 expression: *Krt16*/K16, responding to early postnatal mechanical stress. We have previously demonstrated that *Krt16*^-/-^ mice fail to robustly express *Krt9*/K9 during postnatal development, although the mechanism remained undefined ^17^. Here, we observed a transient rise and fall of *Krt16*/K16 prior to the robust increase of *Krt9*/K9 in normal developing palmoplantar epidermis; however, this elevation of *Krt16*/K16 persists and worsens in *Krt9^-/-^* mice. *Krt16*/K16 likely provides an acute, YAP1-responsive signal that is followed by mechanically sensitive induction of *Krt9*/K9 during the establishment of normal palmoplantar homeostasis. This has additional relevance in disease contexts; as we report here, YAP1 inhibition directly reduces the abnormally high *Krt16*/K16 expression in footpad skin lesions of *Krt9*^-/-^ mice. Previous work has indicated the *Krt16*/K16 is also elevated in the *Krt10^-/-^/Krt2^-/-^*double-null mice, which also experience compensatory *Krt9* upregulation ^70^. Our data suggest that while *Krt16*/K16 is directly *responsive* to YAP1, it is not intrinsically competent to modulate its subcellular distribution and, consequently, its transcriptional activity. Instead, *Krt9*/K9 expression is required to inactivate YAP1 in the palmoplantar epidermis, dampening *Krt16*/K16 expression and mediating the tissue’s adaptation to the experience of mechanical stress. There is, to our knowledge, no precedent for such interdependency between two keratin proteins.

YAP1 signaling is a crucial regulator of mechanotransduction ^18^, and its role in epidermal homeostasis is well-established ^21, 71^. Consistent with previous literature, we find that YAP1 occurs at low levels in the nucleus of differentiating keratinocytes ^22, 23, 71, 72^, while it prominently localizes to the nucleus in keratinocytes within the basal layer ^73, 74^. Our previous work has identified keratins as crucial regulators of this phenomenon; we have proposed a “gating” effect in the progenitor compartment that depends on the occurrence of keratin species with conserved stutter cysteines in the latter portion of the Coil 2 domain ^48^. Our original model, based on studies focused on K14, argues that disulfide-bonded, K14-containing keratin filaments bind 14-3-3σ in a phosphorylation-dependent manner and sequester of YAP1 in the cytoplasm, after which keratinocyte differentiation can proceed. As keratinocytes differentiate, however, K14 ceases to be made, and K10 and, in the palmoplantar epidermis, K9, become the predominant type I keratin species featuring stutter cysteine residues. We therefore predicted the involvement of a “handoff” mechanism whereby cytoplasmic sequestration of YAP1 would necessitate, as differentiation proceeds, the involvement of a differentiation-specific keratin that features the stutter cysteine and can bind 14-3-3 isoforms ^48^. Interestingly, K16 is differentiation-specific, contains a stutter cysteine (**Suppl. Fig. 5A**), and interacts with 14-3-3 proteins ^30^; however, it is not competent to sequester YAP1 (**Fig. 5A**), and its strong expression in *KRT9*-pEDD (**Suppl. Fig. 1**) and *Krt9*^-/-^ plantar skin ^12^ does not rescue the strong nuclear YAP1 localization resulting from K9 loss. The findings reported here provide compelling evidence that K9 fulfils the specialized role of YAP1 sequestration in the epidermis of palmoplantar skin. Importantly, *KRT9*/K9 is the only keratin gene in which a pathogenic variant altering the stutter cysteine has been documented so far ^47^, as the *KRT9* C406→R variant has been shown to be causative for a classical presentation of *KRT9*-pEDD. The data we report here suggest that, in addition to potentially causing fragility of the keratin cytoskeleton ^10, 75^, *KRT9*-pEDD-causative pathogenic variants in *KRT9* crucially fail to sequester YAP1 during differentiation.

A revised and expanded model that incorporates recent findings on K15 (see ^76^) and the current findings on the differentiation-specific K9 is presented in **Supplementary Figure 8**. The model proposes that expression of K15 at sufficiently high levels promotes the progenitor state in the relevant subset of keratinocytes in the basal layer; that the progressive buildup in K14 levels overcomes the influence of K15 and promotes the entry of progenitor keratinocytes into differentiation once specified through post-translational modifications; and that either K9 (palmoplantar epidermis) or K10 (interfollicular epidermis) is required to maintain keratinocytes into the differentiation program (**Suppl. Fig. 8**). In circumstances of high mechanical strain, such as in the palmoplantar epidermis, YAP1-induced K16 in the suprabasal layer provides a signal engaging the expression of *Krt9*/K9 (**Suppl. Fig. 8**). The sequential expression of specific type I keratins in the epidermis (first K15; then K14, then K9 or K10) therefore has a profound impact on the regulation of the balance between the progenitor and differentiating status of keratinocytes and their normal progression through terminal differentiation. Importantly, our model does not propose that keratins regulate proliferation or differentiation *per se*– instead, the expression of specific type 1 keratins promotes cellular states along the homeostatic continuum by modulating effectors of progenitor identity, such as YAP1 (ref. ^76^). While the model relies on the ability of these keratins to sequester YAP1 in the cytoplasm, open questions remain. These include the precise mechanisms through which keratins engage 14-3-3 proteins (which are bivalent adaptor proteins that can function as heterodimers; ^50, 77^) and the role of site-specific phosphorylation on keratins, which is typically required for 14-3-3 interaction ^50, 55^.

The relationship between K9 and YAP1 in the palmoplantar epidermis has significance beyond disease resulting from mutations in *KRT9* itself. Some diseases that manifest with PPK correlate to nonfunctional or low levels of *KRT9*/K9; most notably, a subset of individuals with pachyonychia congenita (PC,^16^) have low levels of K9 in the lesional palmoplantar epidermis. Other palmoplantar epidermal differentiation disorders, such as those arising from variants in desmosome genes, interrupt the mechanical linkage of the keratin filament network to the periphery of the cell ^78, 79^. Additionally, individuals with *WNT10A* variants ^43^ develop PPK due to defects in KLF4 and the subsequent failure to robustly express *KRT9*/K9, and patients with variants directly in KLF4 also develop PPK ^80^. Given the findings described here, it is possible that diverse genetic causes of PPKs converge on the etiology of failed K9 expression, leading to dysregulated YAP1. A survey of *KRT9* expression and YAP1 localization and activity in various palmoplantar epidermal differentiation disorders may allow pharmacological inhibition of YAP1 to be a promising intervention for a subset of PPKs.

Our findings identify K9 as an important modulator of YAP1 signaling in the palmoplantar epidermis, dynamically expressed in response to the mechanical stress borne at this specialized epidermal site. More broadly, K9 joins a growing list of keratin proteins that modulate aspects of keratinocyte differentiation and epidermal homeostasis, especially in the unique context of mechanical stress.

## MATERIALS AND METHODS

### Human sample collection

Collection of healthy human plantar and trunk skin was approved by the University of Michigan Institutional Review Board (IRB HUM00174864), and all participants provided written informed consent. Eligibility criteria required clinically normal, healthy skin from individuals without a history or diagnosis of inflammatory skin disease. Collection of *KRT9*-pEDD patient plantar keratodermas was approved by the Tel Aviv Sourasky Medical Center Institutional Review Board (IRB TLV-0536-08) and National Health Service Health Research Authority London – South East (REC reference 08/H1102/73). Patients presenting with palmoplantar keratodermas were genotyped for variants in *KRT9*. For both healthy volunteers and *KRT9*-pEDD patients, four-millimeter punch biopsies were obtained from acral and healthy neutral (trunk) sites, and subsequently either cryo-embedded or fixed overnight in 10% neutral-buffered formalin, transferred to 70% ethanol, and paraffin-embedded prior to sectioning for immunofluorescence.

### Mouse models, crosses, and treatments

All mouse experiments involved animals in the C57/Bl6J background and were approved by the Institutional Animal Use and Care Committee of University of Michigan Medical School. The *Krt9*^−/−^ [^12^], *Krt16*^-/-^ [^81^], and TEADi [^20^] mouse strains were previously described. Animals were collected at E18.5, P0, P1, P3, P10, and 3.5wks of age. Equivalent numbers of male and female mice were collected. For topical treatment of verteporfin, verteporfin (Sigma-Aldrich, SML0534-25MG) was dissolved at 30mg/mL in 10μl DMSO; immediately prior to use, this stock was diluted to 3mg/mL ^(58,82)^, and 10μl was applied to the ventral surface of the right paw; 10µL vehicle (DMSO) was applied to the left paw. The procedure was repeated once a day for seven days, and animals were collected 24 hours after the final topical application.

TEADi mice were used for the following experiments; these mice carry a tetracycline-inducible synthetic TEADi gene (abbreviated *tetO*-TEADi), conditionally express reversible tetracycline inducible transactivator under the ROSA26 promoter (abbreviated ROSA26 lox-stop-lox [LSL]-rtTA mice), and carry a tamoxifen inducible Cre-recombinase under control of the keratin 14 promoter (K14^CreERT^). For use of the TEADi mice, experimental and control animals were fed with 625mg/kg doxycycline chow (Inotiv, TD.01306) starting 24 hours before first treatment. Tamoxifen (Thermofisher Scientific, #J63509.ME) was dissolved at 2 mg/mL in DMSO and aliquoted in 10µL increments; immediately prior to use, one aliquot per mouse was thawed and applied to the ventral surface of the right paw, while vehicle (DMSO) was applied to the left. The procedure was repeated once a day for 3 days while animals were continually administered doxycycline; paws were collected 7 days after the first treatment (Suppl. Fig. 6). For prenatal TEADi induction, timed pregnant WT dams crossed with TEADi+ (K14^CreERT^ ^+/+^; ROSA26 *^LSL^*^-rtTA^ ^+/+^; *tetO*-TEADi ^+/-^) males. Pregnant dams were fed with 625mg/kg doxycycline chow (Inotiv, TD.01306) starting 24 hours before first treatment. At E18.5, TEADi expression was induced with a single or intraperitoneal injection of 60mg tamoxifen per kg bodyweight (Thermofisher Scientific, #J63509.ME) dissolved in corn oil.

### Tissue collection and cryosectioning

Mice were euthanized and paw tissue was collected at specific time points as indicated. Paw samples were embedded in −40°C optimal cutting temperature compound (O.C.T, Sakura Finetek USA, #4583). Cryosectioning was performed at −20°C using a CRYOSTAR NX50 Cryostat (Thermo Scientific) and MX35 ultramicrotome blade (Epredia #3053835), and 5-μm thick cross-sections were cut and placed on positively charged microscope slides (VWR #48311–703) and stored at −40°C until further use.

### Cell lines and treatments

HeLa and A431 cells were purchased from ATCC and routinely tested for mycoplasma using the MycoAlert® Mycoplasma Detection Kit (Lonza, LT07-118). Cells were cultured in DMEM medium (Gibco #11995–065) supplemented with 10% FBS and 0.01% Penicillin-Streptomycin (Gibco #15140–122). N/TERT keratinocytes (N/TERT-2G) were grown in Keratinocyte SFM media, supplemented with epidermal growth factor (rEGF, 0.2ng/ml), bovine pituitary extract (BPE, 30 ug/ml), calcium chloride (0.31mM) and penicillin streptomycin (10 units/ml). N/TERT cells were seeded at 300,000 (100mm plate) or at 30,000 cells (6 well plate), with media replaced every 48 h. Cells were monitored daily for confluence and monitored for formation of bilayer. HeLa cells were treated with 0-5 μM verteporfin (Sigma-Aldrich, SML0534-25MG) or 3µM VT107 (MedChemExpress #HY-134957) in DMSO for 24 hours post-transfection. N/TERT and A431 cells subjected to compression were treated with 2.5µM verteporfin (Sigma-Aldrich, SML0534-25MG) in DMSO or DMSO vehicle for the entire 24 hours of compression.

### Compression Device Use

N/TERT and A431 cells were cultured as described above. Cells were collected from plates using 0.05% trypsin and pelleted before suspension at 10 million cells/mL in the interpenetrating hydrogel comprising of 1% agarose and 1mg/mL collagen type I (R&D Systems, #3443-100-01), as previously described [^34^]. The cell laden hydrogels were plated within the control or experimental wells of the compression bioreactor and allowed to gel for 15 minutes. Finally, the entire device was placed within the cell culture incubator (5% CO_2_, 37 °C) for 24 hours. Continuous air pressure was supplied by a syringe pump (New Era Infusion One, #NE-300) and monitored for a gauge pressure of 14-15kPa, as extrapolated from COMSOL models (Suppl. Fig. 4).

### COMSOL Computational Modeling

Hydrogel characteristics previously determined through SEM, porosimetery, and rheometry [^34^] were used as inputs for the COMSOL Multiphysics 5.5 computational analysis of applied solid mechanics and are provided in Suppl. Fig. 2. Linear elastic models were used to describe the application force and resultant third principle stress component. Linear elastic properties were assigned to both the hydrogel and membrane material characteristics and the application of pressure was described using an underlying boundary load to define the pressure application on the membrane (Suppl. Fig. 4).

### Indirect immunofluorescence of mouse and human skin

For FFPE tissue sections, slides were cleared using HistoClear (CATALOGUE) for 20 min and rehydrated in progressive 3 minute washes of 100%, 95%, and 70% ethanol, followed by 5 minutes in running water. A circle was drawn around each tissue section with a hydrophobic barrier pen (CALIBIOCHEM #402176), and tissues were blocked with FFPE blocking buffer (10% normal donkey serum, 2% bovine serum albumin in 1X PBS) for one hour at room temperature. For frozen tissue sections, slides stored at −40°C were taken to room temperature, dried for 10 min, fixed with 4% paraformaldehyde (PFA) (Electron Microscopy Sciences #15710) for 20 min at room temperature, followed by 3 washes of 5 min with 1× PBS. A circle was drawn around each tissue section with a hydrophobic barrier pen (CALIBIOCHEM #402176). Tissue sections were blocked with blocking buffer (2% normal donkey serum, 1% bovine serum albumin in 1× PBS) for 1 hour at room temperature. For cultured cell samples, cells were seeded on glass coverslips in 12-well plates and cultured overnight at 37°C and 5% CO2 to let cells attach. After treatment, cells were fixed with 4% PFA for 10 min, permeabilized with 0.5% Triton X-100 for 5 min, then blocked with blocking buffer (5% normal donkey serum in 1× PBS) for 1 h at room temperature. For all conditions, unconjugated primary antibodies (cf. S1 Table) were diluted in blocking buffer(s) and applied overnight at 4°C. On the second day, samples were washed and incubated with fluorophore-conjugated secondary antibodies diluted in blocking buffer(s) for 1 h at room temperature in dark. Samples were then stained with 1 μg/ml of DAPI (Milipore Sigma #268298), washed, mounted with coverslips via FluoroSave reagent (EMD Millipore #345789), and dried overnight. Both tissue sections and cultured cell samples were imaged using a Zeiss LSM800 confocal microscope (Zeiss, Germany). The antibodies used are listed in Table S1. Laser intensity and detector gain were optimized for each fluor/channel. For image quantification, thresholds for positive staining were determined on the basis of fluorescence intensity in control samples.

### Quantification of YAP1 nuclear to cytoplasmic ratio

Cells were stained for YAP1 and imaged as described above. Microscopy images were processed using ImageJ. Using the ImageJ freeform tool, the outer perimeter of the cell body was defined using the GFP channel in cells transfected with GFP-labeled constructs, and the YAP1 intensity mean value was quantified (whole-cell MFI). Next, the nucleus was defined using the DAPI channel and the YAP1 mean intensity value was calculated (nuclear MFI). Nuclear YAP1 MFI was subtracted from whole-cell MFI to generate cytoplasmic MFI. The nuclear-to-cytoplasmic ratio was calculated by dividing nuclear MFI by cytoplasmic MFI. All conditions were normalized relative to the nuclear-to-cytoplasmic ratio of cells transfected with pMAX-GFP.

### Protein Digestion and TMT labeling

Tissue lysates were prepared using TriZOL reagent (Fisher Scientific #15596018) according to manufacturer’s instructions. Protein pellets were solubilized in 8M urea lysis buffer, as previously described ^(17)^. Proteins were denatured by boiling with Laemmli buffer (Bio-Rad #1610747) supplemented with 10% β-mercaptoethanol (β-ME) at 95°C for 10 min. The samples were separated by SDS-PAGE and stained with SimplyBlue SafeStain (Thermo Fisher Scientific). The protein samples were processed and analyzed at the Mass Spectrometry Facility of the Department of Pathology at the University of Michigan. Gel slice(s) corresponding to between 40kDa and 100kDa were taken and destained with 30% methanol for 4 h. Upon reduction (10 mM DTT) and alklylation (65 mM 2-Chloroacetamide) of the cysteines, proteins were digested overnight with 500 ng of sequencing grade, modified trypsin (Promega) at 37° C. Peptides were extracted by incubating the gel with 150 µL of 50% acetonitrile/0.1% TFA for 30 min at room temperature. A second extraction with 150 µL of 100% acetonitrile/0.1% TFA was also performed. Both extracts were combined and dried using a vacufuge (Eppendorf). After reconstitution in 100 μL of 100 mM triethylammonium bicarbonate, the peptides were labeled with a TMT 10plex reagent (ThermoFisher; Cat #90110) following the manufacturer’s protocol. Labeling was quenched by the addition of 8 µL of 5% hydroxylamine. Finally, samples were combined and dried completely before desalting them with SepPak C18 cartridges. The final samples were reconstituted in 20 μL of a 0.1% formic acid/2% acetonitrile solution. Two μL of this solution to collect high-resolution LC-MS (liquid chromatography-mass spectrometry) data in both MS2 and multinotch MS3 modes.

### Liquid chromatography-mass spectrometry analysis (LC-multinotch MS3)

To obtain superior quantitation accuracy, we employed multinotch-MS3 ^(83)^ which minimizes the reporter ion ratio distortion resulting from fragmentation of co-isolated peptides during MS analysis. Orbitrap Ascend Tribrid equipped with FAIMS source (Thermo Fisher Scientific) and Vanquish Neo UHPLC were used to acquire the data. Two µL of the sample was resolved on an Easy-Spray PepMap Neo column (75 µm i.d. x 50 cm; Thermo Scientific) at the flow-rate of 300 nL/min using 0.1% formic acid/acetonitrile gradient system (3-19% acetonitrile in 72 min; 19--29% acetonitrile in 28 min; 29-41% in 20 min, followed by a 10 min column wash using 95% acetonitrile and re-equilibration) and directly sprayed onto the mass spectrometer using EasySpray source (Thermo Fisher Scientific). FAIMS source was operated in standard resolution mode, with a nitrogen gas flow of 4.2 L/min, and inner and outer electrode temperature of 100 °C and dispersion voltage or −5000 V. Two compensation voltages (CVs) of −45 and −65 V, 1.5 seconds per CV, were employed to select ions that enter the mass spectrometer for MS1 scan and MS/MS cycles. Mass spectrometer was set to collect MS1 scan (Orbitrap; 400-1600 m/z; 120K resolution; AGC target of 100%; max IT in Auto) following which precursor ions with charge states of 2-6 were isolated by quadrupole mass filter at 0.7 m/z width and fragmented by collision induced dissociation in ion trap (NCE 30%; normalized AGC target of 100%; max IT 35 ms). For multinotch-MS3, top 10 precursors from each MS2 were fragmented by HCD followed by Orbitrap analysis (NCE 55; 45K resolution; normalized AGC target of 200%; max IT 200 ms, 100-500 m/z scan range).

### Transient transfection of EGFP-tagged keratin constructs

The EGFP-K14WT and EGFP-K16WT constructs (pC3-EGFP vector backbone) have been described (^33, 84^), as have the WT K5 and WTK6-Myc constructs. pMAX-GFP was supplied by Lonza. The K9-GFP and K1-FLAG constructs were purchased from OriGene and GenScript, respectively (Origene Technologies #RG218091, GenScript #OHu19449). The K9 mutant constructs (K9 R163Q/K9 C406A) were generated by site-directed mutagenesis by the U-M Vector Core. Constructs were transiently transfected into HeLa using the SE Cell Line 4D-Nucleofector X Kit (Lonza #V4XC-1032) and Lonza 4D-nucleofector X unit, pulse code CN-114. A431 cells using the SF Cell Line 4D-Nucleofector X Kit (Lonza #V4XC-2032) and Lonza 4D-nucleofector X unit pulse code EQ-100. 1 μg of total plasmid was used to transfect every 0.4 million cells. Transfected cells were plated on coverslips for immunofluorescence assays or in 12-well plates for RNA or protein extraction and RT-qPCR, IP, or WB. . HeLa cells were allowed to rest for 24 h and then subjected to the desired analyses. A431 cells were allowed to rest for 48 hours before harvesting for analyses.

### Proximity ligation assay

A431 cells were transfected using SF Cell Line 4D-Nucleofector kit (Lonza, V4XC-2032) and program EQ-100. 150,000 cells and 1 ug of total plasmid DNA were transfected per parameter. After transfection, cells were plated on #1.5 glass coverslips and cultured for 24 hours. After 24 hours, media was removed, and cells were rinsed with 1X PBS and fixed in 4% PFA/PBS for 10 minutes. Fixed cells were washed, permeabilized for 10 minutes in 0.1% Triton X-100 and blocked in 2.5% NDS/PBS overnight at 4C. Cells were then incubated for 1 hour at 37C with mouse anti-GFP antibody (Novus Biologicals, NB600-597SS) and rabbit anti-14-3-3σ antibody (Sigma-Aldrich, PLA0201). Following primary antibody incubation, cells were incubated with anti-rabbit PLUS and anti-goat MINUS DuoLink Probes (Sigma-Aldrich, DUO92002, DUO92006), and PLA signal was developed according to manufacturer protocol (Sigma-Aldrich, DUO92013).

Coverslips were imaged with a 40X objective with a Zeiss LSM 800 confocal microscope. Laser intensity and detector gain were optimized for each fluor/channel. Images were taken as Z-stacks spanning 10µm at 1µm intervals. ImageJ was used to generate maximum intensity projection images, and the number of PLA punctae per cell was quantified using the ImageJ multi-point tool, using GFP autofluorescence to define the boundaries of the cell(s). Statistical analysis was performed using GraphPad Prism and Mann-Whitney tests.

### Quantitative real-time PCR analysis (qRT-PCR)

For tissue, RNA was isolated using TriZOL (Fisher Scientific, #15596018) according to manufacturer’s protocol. For cells, RNA was isolated using Qiagen RNeasy mini kit (Qiagen #74104) following the manufacturer’s protocol. Total RNA from either tissue or cells was converted to complementary DNA (cDNA) using iScript cDNA Synthesis Kit (Bio-Rad #1708891). The cDNA obtained was subjected to qRT-PCR using the itaq Universal SYBR green kit (Bio-Rad, #1725122) and the CFX 96 Real-Time System (Bio-Rad). The PCR parameters for qRT-PCR were 95°C for 5 min, followed by 40 cycles of 95°C for 10 s and 60°C for 30 s. A “no cDNA template control” and a “melt curve” were included in every PCR run. The normalized expression value of the target gene was determined by first averaging the relative expression of the target gene for each cDNA sample (ΔCq = average *Cq_target_* gene—average *Cq_reference_* gene) and then normalizing the relative expression value of the experimental condition to the control condition (2-(*ΔCq_Experimental_*- *ΔCq_Control_*)). Primers used in qRT-PCR assays are listed in **Table S2**.

### Western blotting and immunoprecipitation

Whole cell lysates were prepared in NP-40 lysis buffer [0.5% NP-40, 150 mM NaCl, 20 mM Tris (pH 7.5), EDTA 1 mM, 1× cOmplete protease inhibitor cocktail solution (Millipore #11836170001), 1× Pierce phosphatase inhibitor cocktail solution (Thermo Scientific #A32957), milliQ water]. Briefly, cultured cells in plates were washed once with cold 1× PBS and transferred to ice. Lysis buffer was added, and cells were scraped off from the plate and rotated at 4°C for 1 h. Supernatants were collected after centrifugation (12,000 × g, 10 min) and total protein levels were measured using Pierce BCA Protein Assay Kits (Thermo Scientific #23227). Proteins were denatured by boiling with Laemmli buffer (Bio-Rad #1610747) containing 10% β-mercaptoethanol (β-ME) for 10 min at 95°C. SDS-PAGE electrophoresis was performed on 4% to 15% gradient gels (Bio-Rad #4561084) or 4% to 20% gradient gels (GenScript #M00656). Gel-bound proteins were transferred to nitrocellulose membranes (Bio-Rad #1620115) using a transblot turbo transfer system (Bio-Rad). Blots were blocked in 5% BSA in phosphate-buffered saline containing 0.1% Tween20 (PBST-T) for 1 h at room temperature and incubated overnight at 4°C with primary antibodies (cf. S1 Table) diluted in the blocking buffer. Secondary antibodies (cf. S1 Table) diluted in blocking buffer were applied for 1 h at room temperature. Blots were developed using SuperSignal West Pico PLUS chemiluminescent substrate (Thermo Scientific #34580) or ECL Select Western Blotting Detection Reagent (Cytiva #RPN2235) and imaged using a FluorChem Q system (ProteinSimple). For immunoprecipitation (IP) of GFP and turbo-GFP tagged constructs, GFP-Trap and turboGFP-Trap Magnetic Agarose (Proteintech, #tbtma-20, #gtma-100) were used, and 25 μl of beads were used. HeLa and A431 cells were lysed and diluted according to manufacturer’s direction. Whole cell lysates (600 μg of total protein) were incubated with beads at 4°C for 1hr. The eluted IP samples were denatured and subjected to western blotting.

### Bulk RNA sequencing

*Krt9*^-/-^ and WT littermates were collected at postnatal day 0 and 3 (N= 3 mice for each group), and RNA was isolated and purified from whole paws using TriZOL reagent (Fisher Scientific, #1559601) using the manufacturer’s instructions. RNA samples were transferred to an external contractor (Novogene) for RNA quality control, RNA ribo-depletion library preparation, and next-generation sequencing (NovaSeq S4 300 cycle). The contractor supplied FASTQ files were mapped to the GRCm39.vm30 reference using STAR version 2.7.10a and the ENCODE standard options from the version documentation ^(85)^. The reference matching GTF file was used with featureCounts from the Bioconductor R package Rsubread to create the counts matrix. The edgeR Bioconductor package was used to filter low expressing genes using the “filterByExpr” function, calculate log counts per million, and normalization factors using the weighted trimmed mean of M-values (TMM) method. The Limma Biocoductor package, with the precision weights, “voom” approach was used to perform linear models ^(86)^. Differential gene expression was determined using custom thresholds as described for each analysis in the main text and in figure legends.

## STUDY APPROVAL

Punch biopsies of healthy trunk and healthy plantar skin were collected with written informed consent under IRB HUM00174864, approved by the University of Michigan IRB. Punch biopsies of *KRT9*-pEDD patient lesions were collected with written informed consent under Tel Aviv Sourasky Medical Center IRB TLV-0536-08 and National Health Service Health Research Authority London – South East (REC reference 08/H1102/73). All samples were deidentified and study authors did not have access to the relevant HIPAA information.

## DATA AVAILABILITY

All data needed to evaluate the findings in this study are present in the paper and/or the Supplementary Materials. RNAseq primary and processed data generated in this manuscript will be deposited in the GEO database after manuscript acceptance. Plasmids generated in this study are available from the lead contact without restriction. Processed RNAseq data is provided in the Source Data File. Code used to generate Supplementary Fig. 4E will be uploaded. The source data underlying Figs. [2B, 3B-C, 4A-A”, 4C-D, 5A-D, 6B, 6D-E, 6G, 7A-B] and Supplementary Figs. [S3D, S4A-C, S5B, S6B-D, S6A-C, S7C], are provided as a Source Data file.

## Lead Contact

Requests for further information and resources should be directed to and will be fulfilled by the lead contact, Pierre A. Coulombe, Ph.D.

## ACKNOWLEDGEMENTS

Research reported in this publication was supported by the National Cancer Institutes of Health under award number P30CA046592 given the use of Rogel Cancer Center Shared Resource(s) including the Tissue and Molecular Pathology Shared Resource and Proteomics Resource Facility. We also thank the Vector Core Facility at the University of Michigan and acknowledge Novogene for their assistance with the bulk RNA sequencing. The authors are grateful to members of the Coulombe laboratory for advice and support, and to the patients and volunteers who donated skin biopsies.

## AUTHOR CONTRIBUTIONS

This study was designed by SNS and PAC. Data was generated by SNS, EH, JYS, and MLK. Data analysis and interpretation were performed by SNS, EH, MA, CNJ, GM, RIB, and PAC. CNJ, MA, LS, OS, ES, EOT, JK, JEG, EH, and GM provided technical and material support. Figures were designed by SNS, with input from EH and PAC. SNS and PAC prepared the manuscript, with input from all authors.

