## Supplementary material for "Under pressure: Keratin 9 regulates mechanosensitive YAP1 signaling in palmoplantar epidermis": Suppl. Figs. 1-8, Suppl. Tables 1-2

**Steiner et al.**

### EXTENDED DATA

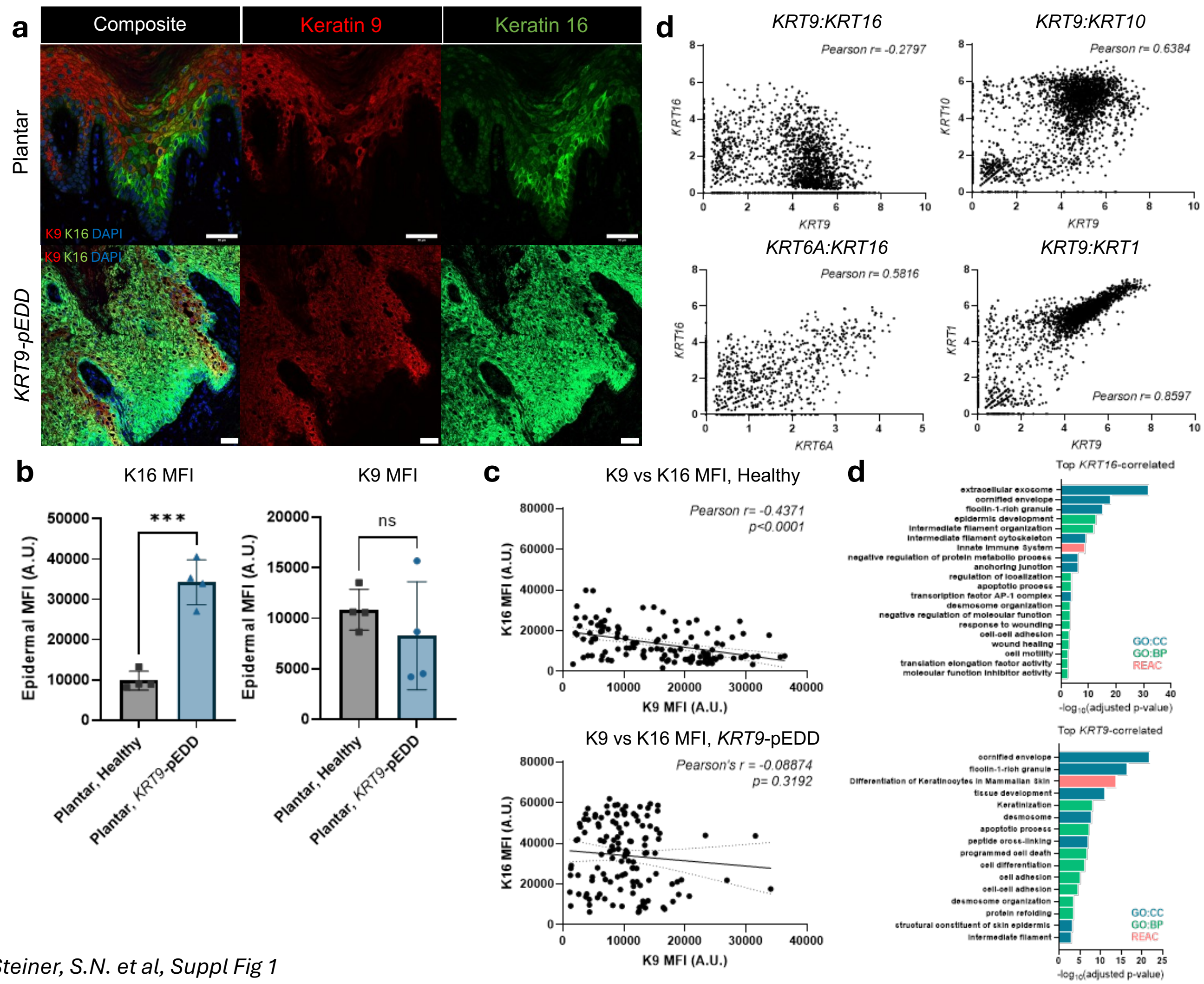

**Supplemental Figure 1: Keratin 16 is elevated and mis-localized in *KRT9*-pEDD epidermis.**

**a**) Immunoflourescence co-staining of K9 and K16 in healthy plantar and *KRT9*-pEDD epidermis. Red= K9, green=K16, blue =DAPI. Scale bar=50μm. **b**) Quantification of K16 and K9 mean fluorescence intensity in healthy plantar epidermis and *KRT9*-pEDD plantar epidermis. Dots represent individuals, with bars representing mean +/- SEM. A.U.= arbitrary units. N=4 healthy and *KRT9*-pEDD individuals. Student t-test analysis. **c**) Quantification of K9 mean fluorescence intensity versus K16 mean fluorescence intensity in single cells in healthy plantar epidermis (top) and *KRT9*-pEDD epidermis (bottom). A.U.= arbitrary units. Healthy epidermis: Pearson r= -0.4371; p<0.0001. *KRT9*-pEDD epidermis: Pearson r= -0.08874, p=0.3192. N=3 healthy and *KRT9*-pEDD individuals counted; approximately 120 cells total counted in each condition. **d**) Transcript pairwise associations palmoplantar keratin(s) in 2,782 cells in single cell RNA datasets from healthy human plantar epidermis. *KRT9* and *KRT16:* Pearson r= -0.28, p<0.0001, *KRT9* and *KRT1:* Pearson r= 0.86, p<0.0001, *KRT9* and *KRT10*: Pearson r= 0.6384, p<0.0001, *KRT16* and *KRT6A:* Pearson r= 0.5816, p<0.0001. **e**) Bar graphs showing all GO categories enriched for *KRT16*-correlated genes (top) and *KRT9*-correlated genes (bottom) with a cutoff of P < 0.05 from GProfiler.

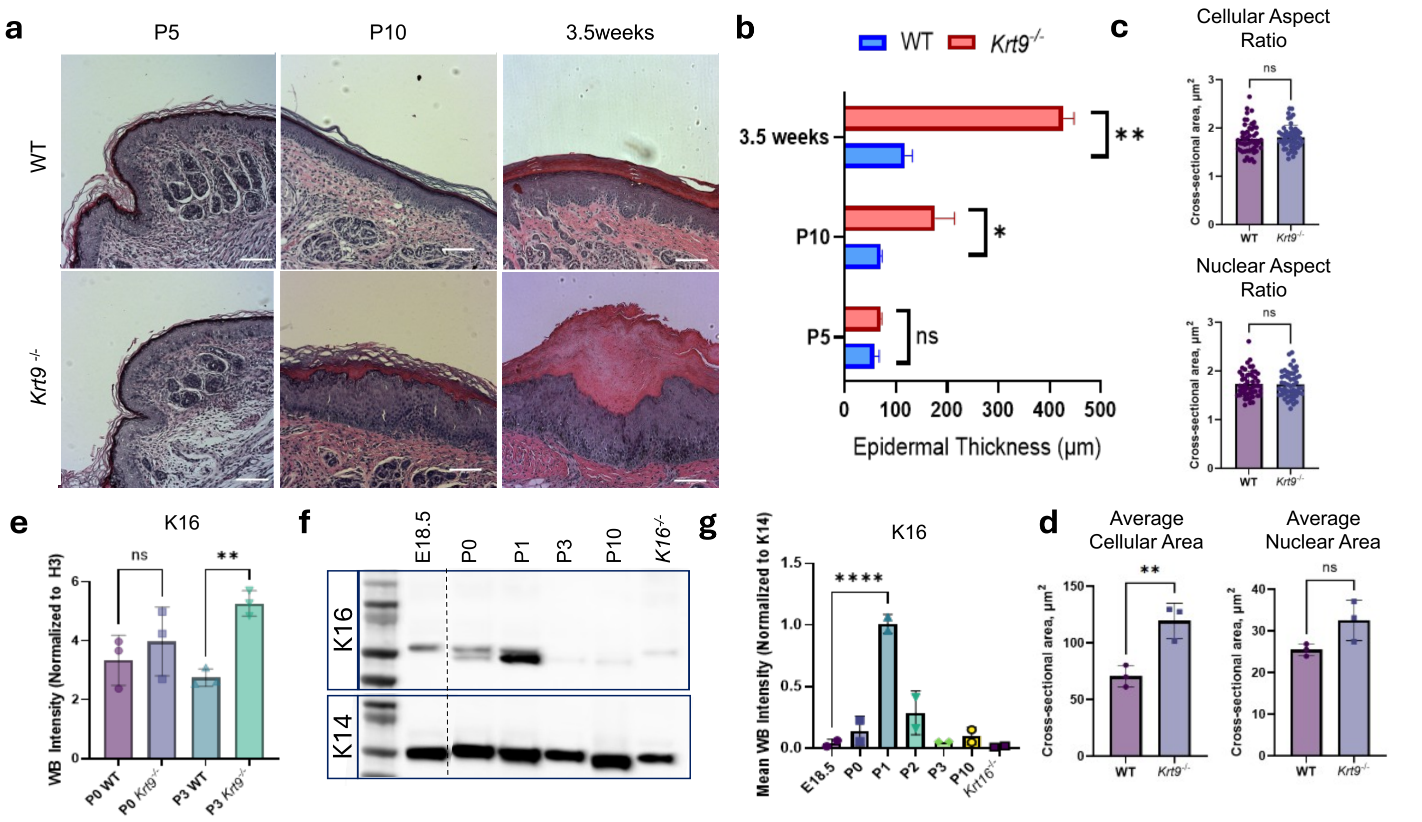

*Steiner, S.N. et al, Suppl Fig 2*

**<u>Supplemental Figure 2</u>: Molecular and histological defects in *Krt9*$^{-/-}$ animals over developmental time.**

**a**) H&E staining of WT/*Krt9*$^{-/-}$ littermates from P5-3.5 weeks of age. N=2 per genotype/timepoint. **b**) Quantification of epidermal thickness. Bars represent mean +/- SEM. Student t-test analysis. **c**) Cross-sectional cellular and nuclear aspect ratio area in palmoplantar epidermis of 3.5-week-old WT/*Krt9*$^{-/-}$ mice. E-cadherin was used to define the edges of the cell, while DAPI was used to define the nucleus. N=3 mice, n=50 cells/mouse quantified. Each dot represents an individual cell. Student t-test. **d**) Cross-sectional cellular and nuclear area in palmoplantar epidermis of 3.5-week-old WT/*Krt9*$^{-/-}$ mice. E-cadherin was used to define the edges of the cell, while DAPI was used to define the nucleus. N=3 mice, n=50 cells/mouse quantified. Each dot represents the average across an individual mouse. Bars represent mean +/- SEM. Student t-test. **e**) Quantification of WB of K16 in P0 and P3 WT/*Krt9*$^{-/-}$ paws. N=3 mice/genotype/timepoint. Each dot represents an individual mouse; bars represent mean +- SEM. One-way ANOVA analysis. **f**) Representative western blot of K16 over WT developmental time. Quantification presented in **g**); N=2 per timepoint. Each dot represents one mouse. Bars represent mean +/- SEM. One-way ANOVA analysis.

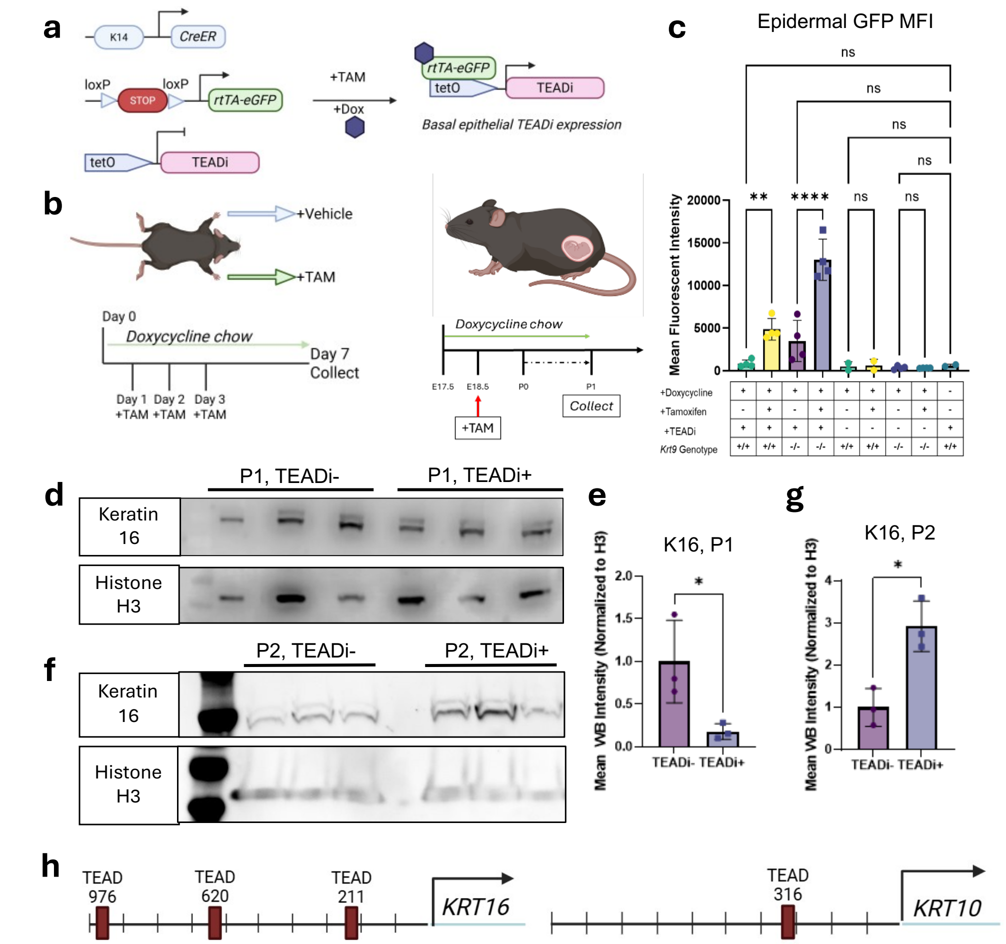

*Steiner, S.N. et al, Suppl Fig 3*

**Supplemental Figure 3: Generation and validation of *Krt9*$^{-/-}$ and WT TEADi+ mouse models.**

**a**) Genetic schematic of the *Krt9*$^{-/-}$ TEADi+ mouse. **b**) Schematic of topical combination doxycycline/tamoxifen treatment schematic of *Krt9*$^{-/-}$ TEADi+ animals and their control counterparts (*Krt9*$^{+/+}$ TEADi+, *Krt9*$^{-/-}$ TEADi-, *Krt9*$^{+/+}$ TEADi-, and TEADi+ animals not administered DOX/TAM), and combination doxycycline/tamoxifen administration in pregnant WT dams bearing E18.5 TEADi +/- pups. **c**) Epidermal GFP autofluorescence measured across all treatment and genotype conditions, demonstrating effective recombination only in TEADi+ samples administered both DOX and TAM. Data represented as mean +/- SEM. N=4 animals/condition. One-way ANOVA. **d**) Western bot of K16 in TEADi+ and TEADi- littermates at postnatal day 1, quantified in **e**). N=3. Each dot represents one mouse. Normalized relative to histone H3; student t-test analysis. **f**) Western blot of K16 in TEADi+ and TEADi- littermates at postnatal day 2, quantified in **g**). N=3. Each dot represents one mouse. Normalized relative to histone H3; student t-test analysis. **h**) Schematic representing location(s) of predicted TEAD motifs upstream of the *KRT16* transcription start site (TSS) and the *KRT10* TSS. Sites identified using the UCSC Genome Browser [1].

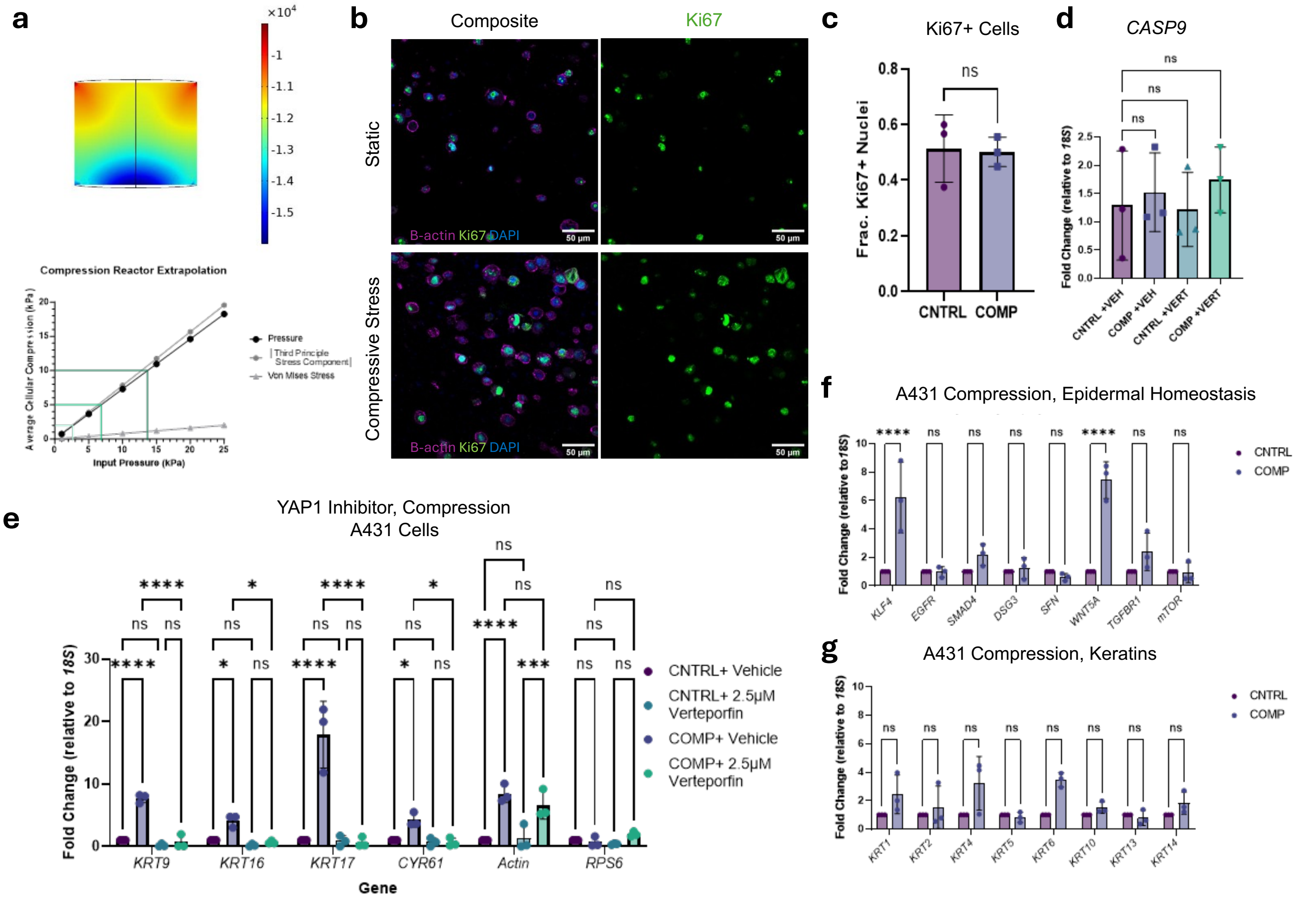

Steiner, S.N. et al, Suppl Fig 4

**<u>Supplemental Figure 4</u>: Compressive force COMSOL modeling and A431 response to compression.**

**a**) Mesh construction on COMSOL model of the hydrogel (1 cm height by 6 mm radius) and deflectable membrane (1 mm height by 6mm radius). COMSOL was used to calculate extrapolation of input air pressure versus modeled internal compressive force using 3 methodologies on 1% agarose/1 mg/mL collagen 1 gel. Simple pressure was determined to be the best model. **b**) IF of Ki67 in control and compressed A431 cells, quantified in **c**). Purple =β-actin, green=Ki67, blue=DAPI. Scale bar= 50μm. N=3. Data represented as mean +/- SEM. Student t-test analysis. **d**) RT-qPCR data of *CASP9* in control and compressed cells, with and without verteporfin (VERT). N=3. Data represented as mean +/- SEM. One-way ANOVA analysis. **e**) A431 cells under compression recapitulate observed results from N-TERT keratinocytes. RT-qPCR of vehicle- or VERT-treated A431 cells under compression of *KRT9*, *KRT16*, *KRT17*, *CYR61, Actin,* and *RPS6*. Dark purple= control -verteporfin, light purple= compressed (10kPa, 24hrs) -verteporfin; dark blue=control +VERT light blue= compressed (10kPa, 24hrs) +VERT. Fold change relative to *18S*. N=3. Two-way ANOVA. Data are represented as mean +/- SEM. **f**) RT-qPCR of a select panel of genes related to epidermal homeostasis and stress response in A431 cells with and without compression. Dark purple=control, dark blue=compressed (10kPa, 24hrs). Fold change relative to *18S*. N=3. Two-way ANOVA. Data are represented as mean +/- SEM. **g**) RT-qPCR of a panel of keratin genes in A431 cells with and without compression. Dark purple=control, dark blue=compressed (10kPa, 24hrs). Fold change relative to *18S*. N=3. Two-way ANOVA. Data are represented as mean +/- SEM.

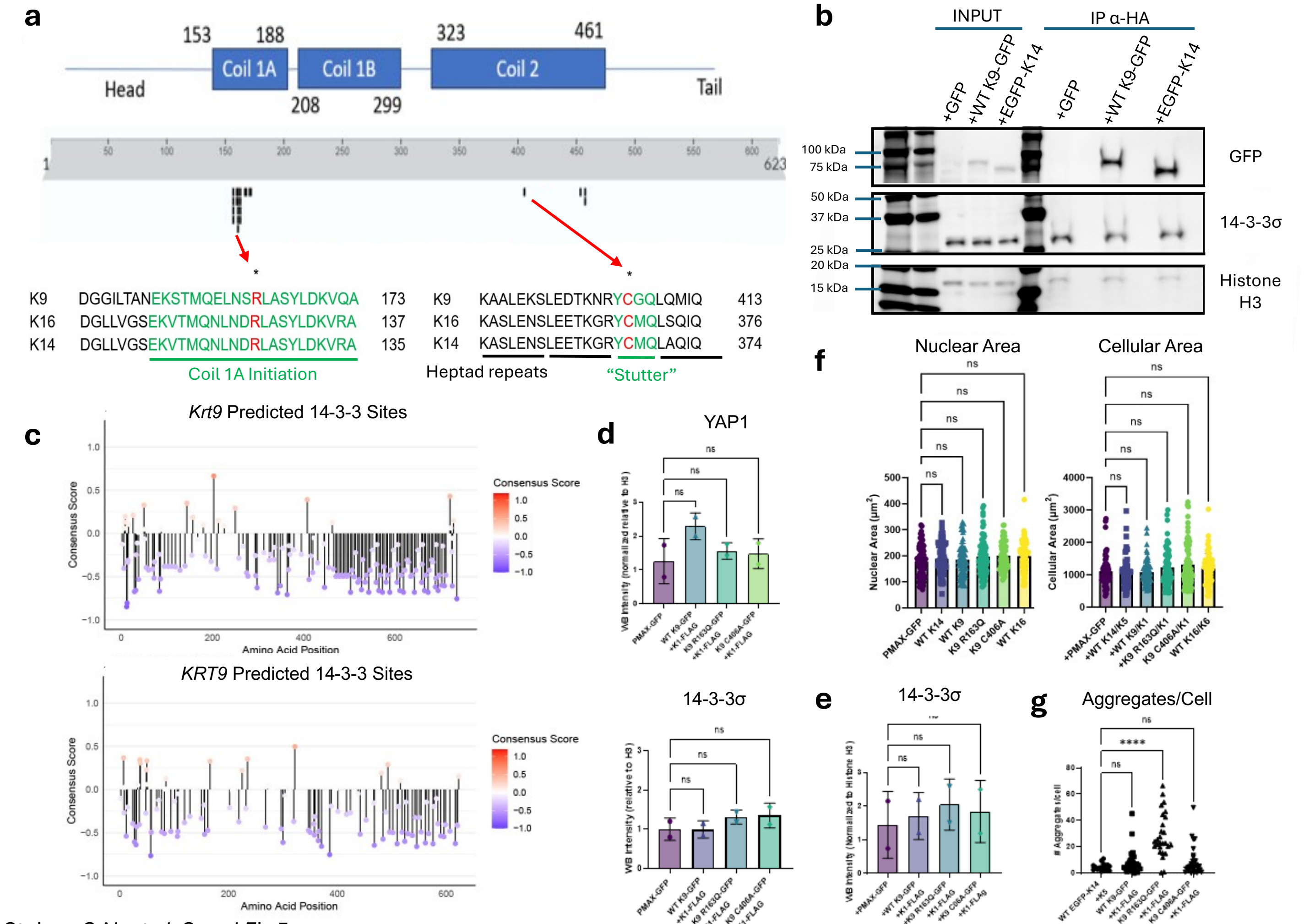

Steiner, S.N. et al, Suppl Fig 5

**<u>Supplemental Figure 5</u>: Analysis of *KRT9*-pEDD-mimic constructs and biochemical phenotypes.**

**a**) Schematic of pathogenic variants in K9 and mutagenized residues; the R163 residue frequently mutated in *KRT9*-pEDD is conserved across K16 and K14, as is the stutter cysteine residue. **b**) Western blot of co-immunoprecipitation of GFP-tagged keratins with HA-tagged 14-3-3σ. IN=input fraction, 10ug total protein loaded. IP=precipitated fraction, 600ug total protein loaded. kDa= kiloDaltons (approx. molecular weight). **c**) Computational prediction of 14-3-3 binding site(s) on mouse and human keratin 9. Red dots represent amino acids highly predicted to be phosphorylated and recognized by 14-3-3 proteins; blue dots represent amino acids predicted to not be bound by 14-3-3 proteins. Sequence via UniProt; analysis performed using 14-3-3-Pred [2]. **d**) Quantification of western blots for D) YAP1 and 14-3-3σ of HeLas transfected with WT and *KRT9*-pEDD mimic constructs. N=2 biological replicates. Data represented as mean +/- SEM. One-way ANOVA analysis. **e**) Western blot quantification of endogenous 14-3-3σ in A431 cells transfected with WT and *KRT9*-pEDD constructs. N=2 biological replicates. One-way ANOVA analysis. **f**) Quantification of nuclear and cellular area in HeLa cells transfected with WT and *KRT9*-pEDD constructs. N=3 biological replicates, n=30 cells per replicate per condition quantified. Each dot represents the nuclear or cellular area of a single cell. One-way ANOVA analysis. **g**) Quantification via R code of punctate GFP aggregates in HeLa cells transfected with WT and *KRT9*-pEDD constructs. N=3, minimum 50 cells/condition/replicate. Dots represent average number of aggregates per cell in a field of view. Data represented as mean +/- SEM. One-way ANOVA analysis.

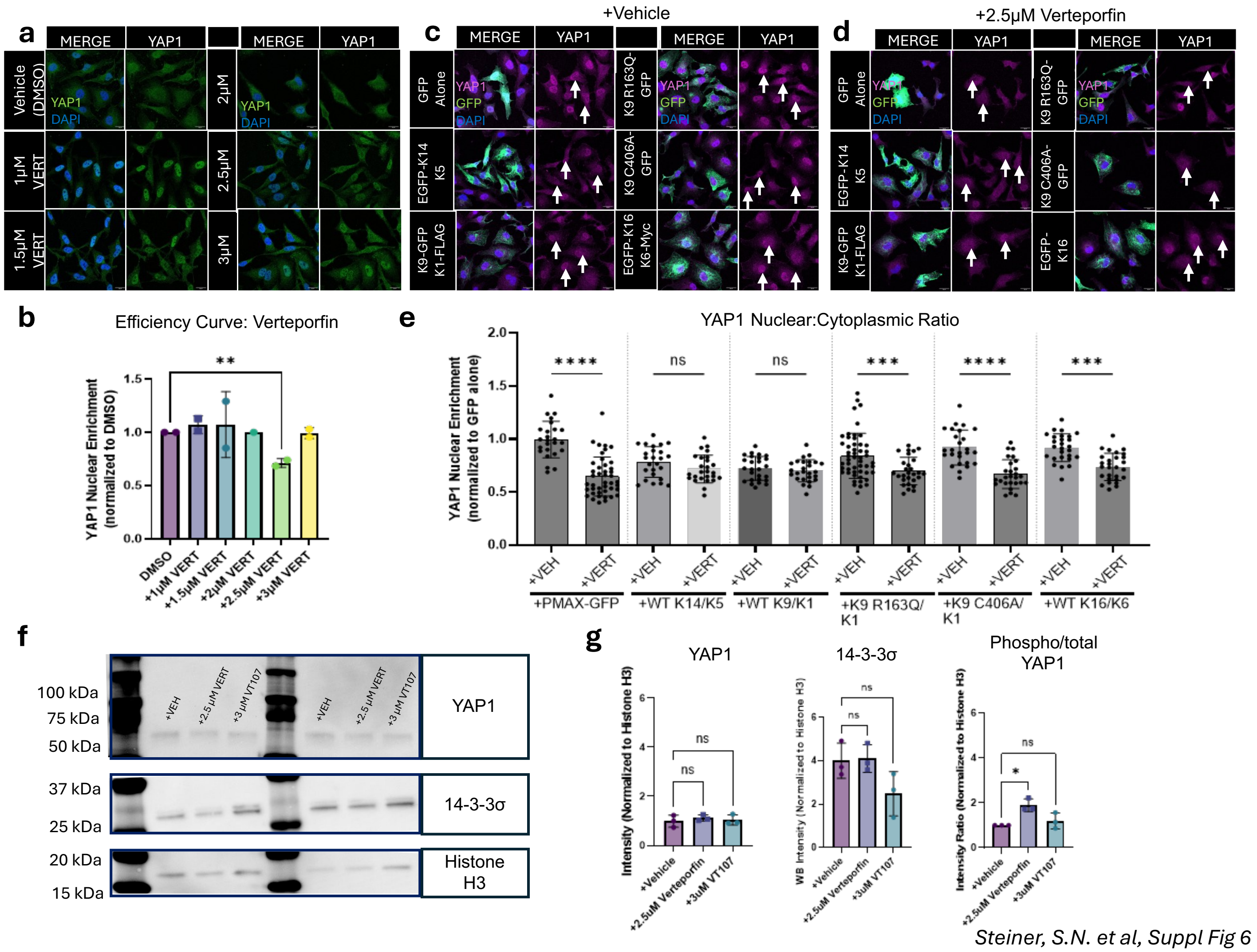

Steiner, S.N. et al, Suppl Fig 6

**<u>Supplemental Figure 6:</u> Verteporfin re-localizes YAP1 to the cytoplasm and reduces its transcriptional activity *in vitro*.**

**a**) IF of YAP1 in HeLas treated with concentrations of verteporfin ranging from 0-5 μM. Blue=DAPI, \ green=YAP1. Scale bar=10μm. **b**) Quantification of (**a**). Average of YAP1 nuclear/cytoplasmic mean intensity, N=2. One-way ANOVA analysis. **c**) IF of HeLa cells transfected with WT and *KRT9*-pEDD GFP-tagged keratin constructs. **d**) IF of HeLa cells transfected with GFP-tagged WT and *KRT9*-pEDD constructs, treated with 2.5μM verteporfin. Blue=DAPI, purple=YAP1, green=GFP-tagged construct autofluorescence. Scale bar =20um. **e**) Quantification of YAP1 fluorescence nuclear to cytoplasmic ratio in (**c-d**). N=3, 50 cells/condition/replicate. Dots represent individual cell value(s), bars representing mean +/- SEM. One-way ANOVA analysis. **f**) WB of 14-3-3σ, YAP1, and Histone H3 in verteporfin-treated HeLa cells. **g**) Quantification of YAP1, 14-3-3σ and phosphorylated-to-to-total YAP1 ratio in verteporfin and VT107-treated HeLa cells. Normalized relative to Histone H3. N=3. Data represented as mean +/- SEM. One-way ANOVA.

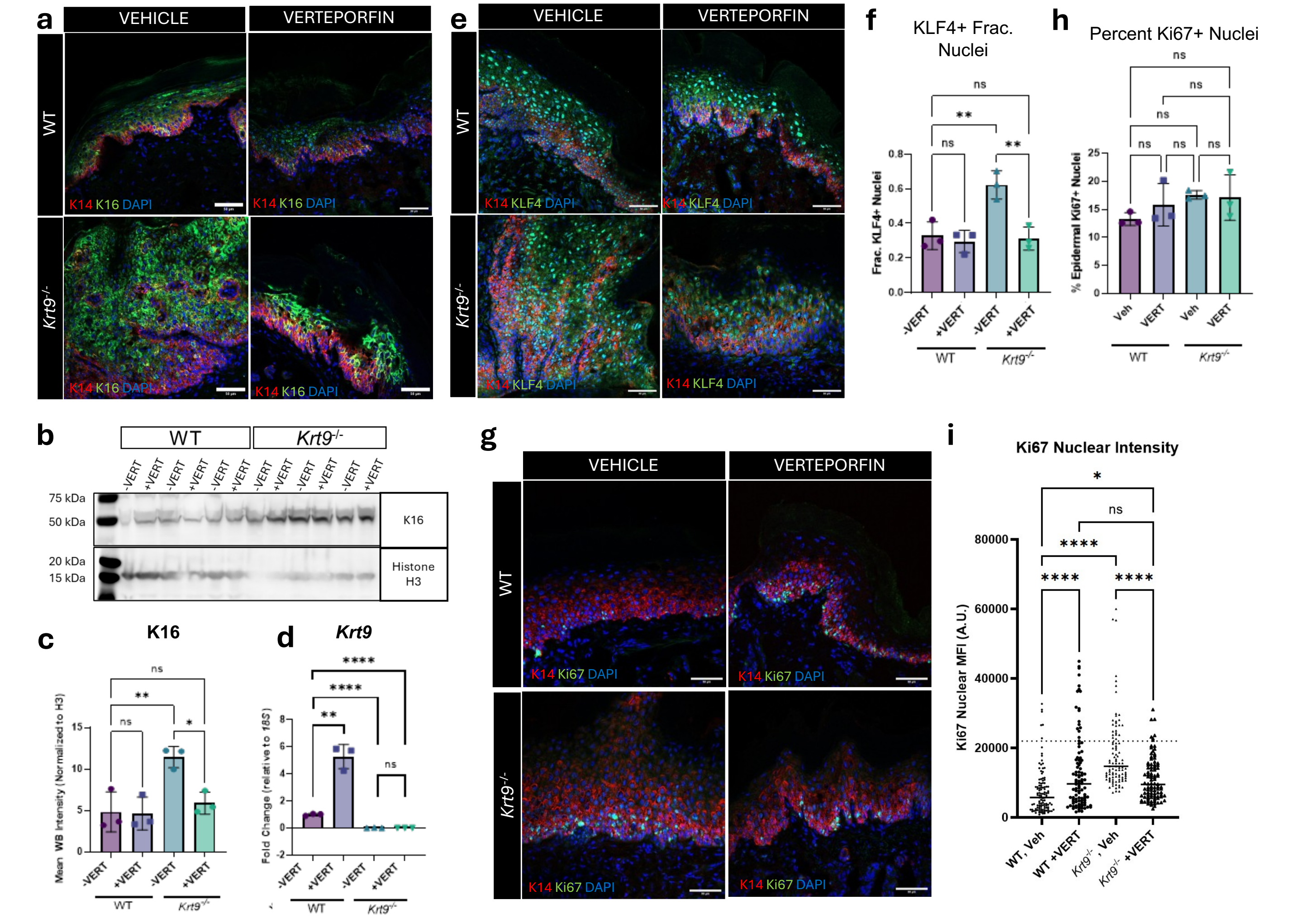

*Steiner, S.N. et al, Suppl Fig 7*

**<u>Supplemental Figure 7</u>: Verteporfin treatment normalizes molecular phenotypes in the *Krt9*$^{-/-}$ mouse.**

**a**) IF of K16 in WT and *Krt9*$^{-/-}$ animals treated with VERT. Red=K14, green=K16, blue=DAPI. Scale bar= 50µm. N=2 animals/condition. **b**) Western blot of K16 and Histone H3 in WT/*Krt9*$^{-/-}$ animals treated with verteporfin. N=3 animals/condition. Data represented as mean +/- SEM. One-way ANOVA. **c**) RT-qPCR of *Krt9* from WT and *Krt9*$^{-/-}$ paws treated with verteporfin. N=3 animals/condition, n=3 technical replicates; dots represent average of 3 technical replicates per individual mouse. Data represented as mean +/- SEM. One-way ANOVA. **d**) RT-qPCR of *Krt9* in WT/*Krt9*$^{-/-}$ mice treated with vehicle or verteporfin (VERT). N=3 animals/condition. Fold change relative to *18s*. Dots represent individual mice; bars represent mean +- SEM. One-way ANOVA analysis. **e**) IF of KLF4 in WT/*Krt9*$^{-/-}$ animals treated with verteporfin. Red=K14, green=KLF4, blue=DAPI. Scale bar=50µm. Quantification of fraction of total nuclei that are KLF4+ in **f**). N=3 animals/condition. Dots represent average KLF4+ fraction in each mouse. Data represented as mean +/- SEM. One-way ANOVA. **g**) IF of Ki67 in WT/*Krt9*$^{-/-}$ animals treated with verteporfin. Red=K14, green=Ki67, blue=DAPI. Scale bar=50µm. Quantification of fraction of total nuclei that are Ki67+ in **h**). N=3 animals/condition. Dots represent average Ki67+ fraction in each mouse. Data represented as mean +/- SEM. One-way ANOVA. **i**) Quantification of nuclear Ki67 intensity in WT and *Krt9*$^{-/-}$ mice treated with vehicle or verteporfin. N=3 mice/condition, n=50 cells counted/mouse. Each dot represents the nuclear Ki67 intensity in an individual cell. Dotted line represents threshold of visually positive staining. A.U.= arbitrary units. One-way ANOVA.

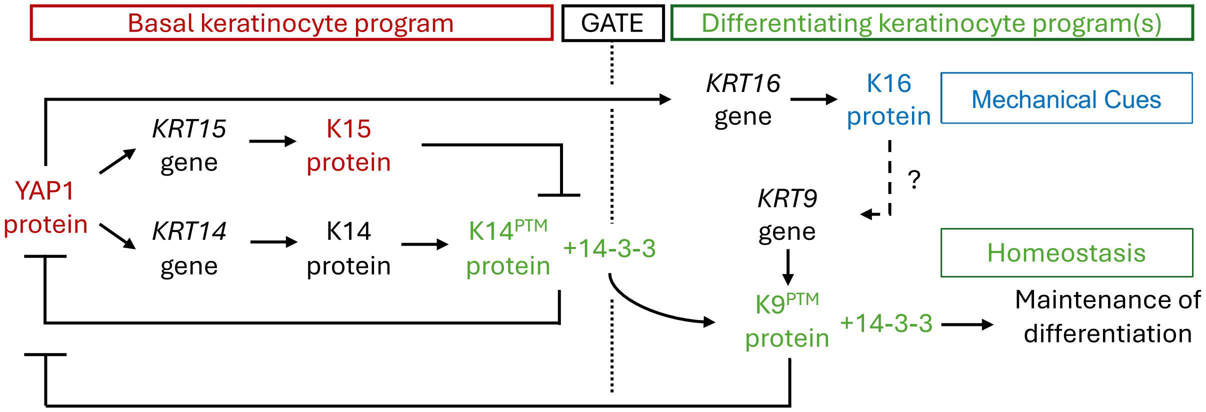

Steiner et al. Supplementary Figure 8

**<u>Supplemental Figure 8</u>: Model.**

Keratin-dependent YAP1 regulation in homeostasis and under stress in the epidermis. See text for details.

### SUPPLEMENTAL TABLES: TITLES AND LEGENDS

#### Table S1: Primary and secondary antibodies used in this study.

| PRIMARY ANTIBODY | | | | | |
|---|---|---|---|---|---|
| Antibody Target | Host species | Catalogue | Manufacturer | Concentration, IF | Concentration, WB |
| Keratin 9 | Guinea pig | GP-CK9 | Progen | 1:500 | N/A |
| Keratin 16, human | Rabbit | - | Takahashi et al, 1994 | 1:500 | N/A |
| YAP1 | Rabbit | 14074S | Cell Signaling | 1:500 | 1:1000 |
| Keratin 16 | Rabbit | - | Bernot and Coulombe, 2002 | 1:500 | 1:1000 |
| Keratin 14 | Chicken | 906004 | BioLegend | 1:500 | 1:1000 |
| E-Cadherin | Mouse IgG2A | 610181 | BD Biosciences | 1:500 | N/A |
| KLF4 | Rabbit | 11880-1-AP | Proteintech | 1:500 | N/A |
| Ki67 | Rabbit | 9129S | Cell Signaling | 1:500 | N/A |
| 14-3-3σ | Rabbit | PLA0201 | Sigma Aldrich | 1:500 | 1:1000 |
| GFP | Mouse | G1546-25UL | Sigma Aldrich | "1:500" | 1:1000 |
| Histone H3 | Rabbit | 4499S | Cell Signaling | N/A | 1:1000 |
| TurboGFP | Rabbit | TA150071 | Origene | N/A | 1:1000 |
| phospho-YAP1 S127 | Rabbit | ab76252 | Abcam | N/A | 1:1000 |
| **SECONDARY ANTIBODY** | | | | | |
| Anitbody Target | Host species | Catalogue | Manufacturer | Concentration, IF | Concentration, WB |
| Anti-guinea pig IgG AF555 | Goat | A-21435 | Invitrogen | 1:500 | N/A |
| Anti-rabbit IgG AF488 | Goat | ab150077 | Abcam | 1:500 | N/A |
| Anti-rabbit IgG AF647 | Goat | A-21244 | Invitrogen | 1:500 | N/A |
| Anti-mouse IgG2A AF555 | Goat | A21137 | Invitrogen | 1:500 | N/A |
| Anti-chicken IgY AF647 | Goat | ab150171 | Abcam | 1:500 | N/A |
| Anti-rabbit IgG HRP | Goat | 7074S | Cell Signaling | N/A | 1:1000 |
| Anti-chicken IgY HRP | Goat | A16054 | Invitrogen | N/A | 1:1000 |
| Anti-mouse IgG HRP | Goat | 7076S | Cell Signaling | N/A | 1:1000 |

#### Table S2: Genotyping and RT-qPCR primers used in this study.

All primers were purchased from IDT. Genotyping primers were designed based on previous publications (ref. [3,4-6]). RT-qPCR and ChIP-qPCR primers were optimized using the IDT PrimerQuest tool.

| Genotyping Primers | | |
|---|---|---|
| Target | Species | Sequence |
| *Krt9* | Mouse | FW WT: GGGATCCCATGCCCTTTTCT |
| | | FW MUT:TCATTCTCAGTATTGTTTTGCC |
| | | RV: TCAGGACAAGGAAGGGAGTG |
| K14-CreERT | Mouse | FW, Internal Control: CTAGGCCACAGAATTGAAAGATCT |

| | | |
|---|---|---|
| | | RV, Internal Control: GTAGGTGGAAATTCTAGCATCC |
| | | FW, Mutant: CGCATCCCTTTCCAATTTAC |
| | | RV, Mutant: GGGTCCATGGTGATACAAGG |
| TEADi | Mouse | FW:CGCGTTAAGTGCAACACGAT |
| | | RV: GAGAAACACTGGACGCCGTA |
| rtTA | Mouse | FW: CTGGCTTCTGAGGACCG |
| | | RV, Mutant: AGACTGCCTTGGGAAAAGCG |
| | | RV, WT: AGCCTGCCCAGAAGACTCC |
| **RT-qPCR Primers** | | |
| Target | Species | Sequence |
| *Krt9* | Mouse | FW: GGTAGCTATGGTGGAGGAAATAG |
| | | RV: GGGTTCTAGTATCGCATCTTGT |
| *KRT9* | Human | FW: GGAGGTGATGGTGGTATTC |
| | | RV: GATAAGGTGCAGGCTCTAG |
| *Krt16* | Mouse | FW: TGAGATGAGGGACCAGTATGA |
| | | RV: TGCGGTTGCTCTGGATTAG |
| *KRT16* | Human | FW: GAGCAGATGGCAGAGAAA |
| | | RV: CTGTACCAGTTCGCTGTT |
| *KRT17* | Human | FW: GGTGGGTGGTGAGATCAATGT |
| | | RV: CGCGGTTCAGTTCCTCTGTC |
| *CYR61* | Human | FW: TTCACTGCTGTATCCCAATAAG |
| | | RV: AGTGTATGCCATTCGGTATTT |
| *Actin* | Human | FW: CATGTACGTTGCCCAGGC |
| | | RV: CTCCTTAATGTCACGCACGAT |
| *18S* | Human | FW: CCTGTGCCTTCCTTGGA |
| | | RV: CATTCGAACGTCTGCCCTATC |
| *Krt14* | Mouse | FW: AGAGTGAGATTTCTGAGC |
| | | RV: GTCTCCAGGTTATTC |
| *18S* | Mouse | FW: CCTGTGCCTTCCTTGGA |
| | | RV: CATTCGAACGTCTGCCCTATC |
| *GAPDH* | Human | FW: GGGGTCATTGATGGCAACAAT |
| | | RV: GGGGTCATTGATGGCAACAATA |
| *RPS6* | Human | FW: TGGACGATGAACGCAAACTTC |
| | | RV: TTCGGACCACATAACCCTTCC |
| **ChIP-qPCR Primers** | | |
| *KRT16* TEAD 211 | Human | FW: GAAAGGGAGCCTGATTC |
| | | RV: ATCAACTCTGCTGTGTG |
| *KRT16* TEAD 620 | Human | FW: AGTGCTGTTTGTATGGT |
| | | RV: ATCCTATAACTCTCTTTCCC |
| *KRT16* TEAD 976 | Human | FW: GTGCCTGTGATTAGGAG |
| | | RV: TCACACTCACCTCTACA |
| *KRT10* TEAD 316 | Human | FW: GCAAACATTCTATGCAACCTGAA |
| | | RV: CACCTGTTGAAGAACATTGAGC |
| *CYR61* TEAD binding site | Human | FW: CACACACACACACACACAAAG |
| | | RV: ATCTCAGGAATGCTGGTTGG |
